# MFF-dependent mitochondrial fission regulates presynaptic release and axon branching by limiting axonal mitochondria size

**DOI:** 10.1101/276691

**Authors:** Tommy L. Lewis, Seok-Kyu Kwon, Annie Lee, Reuben Shaw, Franck Polleux

**Author notes:** These authors contributed equally to this work (Tommy L. Lewis, Seok-Kyu Kwon). Address correspondence to: Franck Polleux Columbia University Department of Neuroscience Mortimer B. Zuckerman Mind Brain Behavior Institute Kavli Institute for Brain Science 1108 NWC Building - MC 4862 550 West 120th Street New York, N.Y. 10027.

## Abstract

Neurons display extreme degrees of polarization, including compartment-specific organelle morphology. In cortical pyramidal neurons, dendritic mitochondria are long and tubular whereas axonal mitochondria display uniformly short length. Here, we explored the functional significance of maintaining small mitochondria for axonal development *in vitro* and *in vivo*. We report that the Drp1 ‘receptor’ Mitochondrial fission factor (MFF) is required for determining the size of mitochondria entering the axon and then for maintenance of their size along the distal portions of the axon without affecting their trafficking properties, presynaptic capture, membrane potential or capacity for ATP production. Strikingly, this increase in presynaptic mitochondrial size upon MFF downregulation augments their capacity for Ca^2+^ ([Ca^2+^]_m_) uptake during neurotransmission, leading to reduced presynaptic [Ca^2+^]_c_ accumulation, decreased presynaptic release and terminal axon branching. Our results uncover a novel mechanism controlling neurotransmitter release and axon branching through fission-dependent regulation of presynaptic mitochondrial size.

## INTRODUCTION

Neurons are among the most highly polarized cells found in nature. This high level of polarization underlies the formation of structural compartments such as the axon, dendrites and spines which in turn dictates information processing and transfer in neural circuits. In order to set up this extreme degree of compartmentalization and maintain it throughout the life of the organism, critical cellular and molecular effectors of polarization, including mRNAs, proteins, and organelles, must be either differentially trafficked or functionally regulated in a spatially precise way ^1^.

Mitochondria are one of the most abundant organelles found in neurons and are localized throughout the axonal and somatodendritic domains ^2–6^. Several studies have observed that dendritic mitochondria have a long and tubular shape, while axonal mitochondria appear to be short and punctate ^2–10^. However, the functional implications of this structural difference have not been addressed. In general, mitochondria play many important physiological functions such as ATP production via oxidative phosphorylation, Ca^2+^ uptake, lipid biogenesis, and can trigger apoptosis through cytochrome-c release ^11^. Dendritic mitochondrial morphology and function have mainly been studied in the context of neurodegenerative diseases and their contribution to activity-dependent homeostasis ^4^,^6^,^12^. In axons, the presynaptic capture of mitochondria is both necessary and sufficient for terminal axon branching in cortical neurons ^13^ and these presynaptically captured mitochondria generate ATP and buffer Ca^2+^ during synaptic transmission ^14–16^. Interestingly, presynaptic Ca^2+^ clearance by individual mitochondria underlies bouton-specific regulation of presynaptic vesicle release along cortical and hippocampal axons ^15,16^. Mitochondrial function has been suggested to differ in specific neuronal compartments ^4,17^, but the outcome of disrupting mitochondrial size in a compartment-specific manner remains to be determined.

Mitochondrial size is regulated through the competing processes of fission and fusion ^18^. Mitochondrial fusion is regulated by two distinct molecular effectors: Mfn1/2 mediates outer membrane fusion ^19,20^ while Opa1 regulates inner membrane fusion ^21,22^. Fission occurs by the ‘pinching’ of a mitochondrion into two new mitochondria via oligomerization of Drp1, a dynamin-like GTPase, at the outer membrane ^23–25^. Because Drp1 is a cytoplasmic protein, it must be recruited to the outer mitochondrial membrane via Drp1 ‘receptors’. There are currently four known Drp1 receptors: MFF, FIS1, MiD49 (Scmr7) and MiD51 (Scmr7L), whose relative contribution to Drp1-dependent fission is still debated ^26–36^.

Our understanding of the relative contribution of fission and fusion for neuronal morphogenesis or synaptic function remains fragmented. Loss of Drp1 is embryonic lethal and leads to defects in brain development, neuronal process outgrowth and synapse formation. However, Drp1 affects many cellular processes other than mitochondrial fission and also severely affects neuronal viability, so this observation does not unequivocally implicate mitochondrial fission as the cause of the defects in neuronal development ^37–42^. Mitochondrial fusion is also required for proper development as each of the Mfn1, Mfn2 and Opa1 knockouts are lethal^20,43,44^, however their role in cortical neuron development is less well studied.

We establish that the small mitochondrial size characterizing the axon of cortical pyramidal neurons is dependent on mitochondrial fission factor (MFF) but not FIS1, the other abundant Drp1 receptor in these neurons. We find that downregulation of *Mff* results in a striking increase of the size of mitochondria entering the axon, and an inversion of the fission/fusion ratio along the axon. This elongation of axonal mitochondria does not affect dendritic mitochondria, presynaptic capture, trafficking along axons, mitochondrial membrane potential or the capacity to generate ATP. However, this Mff-dependent increase in presynaptic mitochondrial length significantly increased total Ca^2+^ uptake capacity, resulting in decreased evoked neurotransmitter release and decreased terminal axon branching *in vivo*. Our results identify a novel molecular mechanism limiting presynaptic mitochondrial size and demonstrate its functional importance for neurotransmitter release and axon branching via the limitation of mitochondrial Ca^2+^ buffering capacity.

## RESULTS

### Mitochondrial Morphology is Strikingly Different in Axons and Dendrites

In order to visualize mitochondrial morphology and function in developing and adult cortical neurons *in vitro* and *in vivo*, we implemented a strategy for sparse labeling via *ex utero* or *in utero* electroporation (EUE and IUE respectively) ^45,46^. We developed Flp recombinase-dependent plasmids expressing either a cytoplasmic fluorescent protein (tdTomato (red), EGFP (green) or mTAGBFP2 (blue)), or a mitochondrial matrix-targeted fluorescent protein (mt-YFP, mt-DsRED), based on a previous report ^47^. When co-electroporated with a low concentration of Flp-e plasmid at E15.5, this strategy resulted in sparse labeling of layer 2/3 cortical pyramidal neurons ^13,48,49^, and allowed us to visualize mitochondrial morphology at high resolution in single, optically-isolated neurons both *in vitro*(with EUE) and *in vivo* (with IUE). Following EUE at E15.5, we acutely dissociated and cultured cortical pyramidal neurons at high density for three weeks (21 Days In Vitro, (DIV)) and observed striking differences in mitochondrial morphology and occupancy between axons and dendrites (**Fig. 1a-c**). In dendrites, mitochondria were elongated (0.52 to 8.88 μm (10-90^th^ percentile)) and occupied the majority of proximal and distal dendritic processes (69.6 ± 2.54%), while throughout the axon, mitochondria are short, relatively standard in length (0.3 to 1.08 μm (10-90^th^ percentile)) and occupied only an average of 4.95 (± 0.4%) of the axonal length. To confirm this was not an *in vitro* artifact, we also performed IUE at E15.5 and visualized sparsely labeled layer 2/3 cortical neurons at (P)ostnatal day 21. Remarkably, quantitative measurement of mitochondrial size and occupancy were highly conserved *in vivo* (**Fig. 1d-f**) where dendritic mitochondria measured 1.31 to 13.28 μm (10-90^th^ percentile) in length and occupied 69.58 ± 2.23% of dendritic processes, while axonal mitochondria measured 0.45 to 1.13 μm (10-90^th^ percentile) and occupied 8.41 ± 0.75% of axon length (**Fig. 1g-h**). These observations confirm that mitochondrial morphology is strikingly different in the axonal and dendritic compartments of cortical layer 2/3 pyramidal neurons both *in vitro* and *in vivo*.

**Figure 1:**
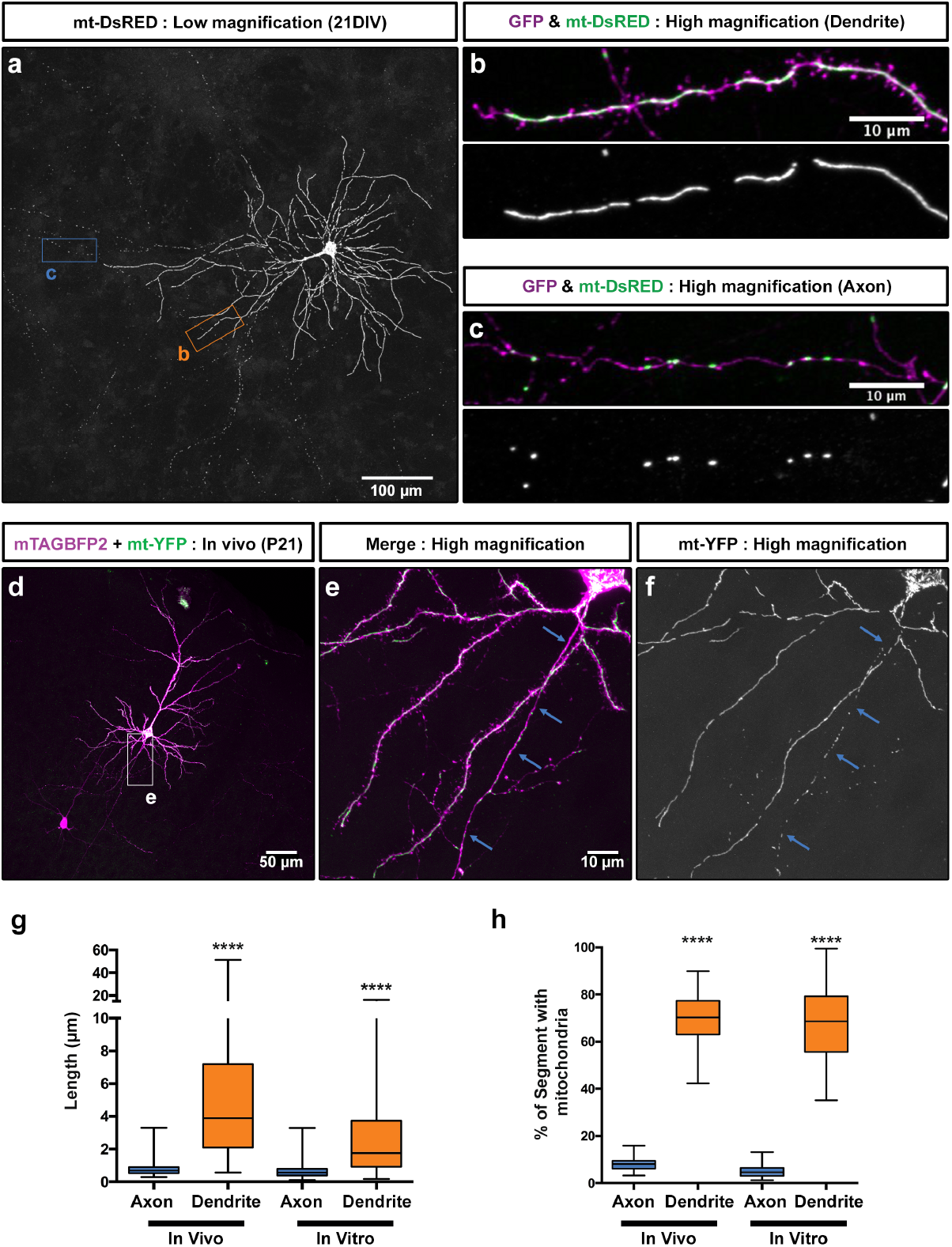
Mitochondria morphology is compartmentally regulated in cortical neurons *in vitro* and *in vivo*. (**a**) Mitochondria morphology visualized by cortical *ex utero* electroporation of a genetically encoded, mitochondria matrix targeted DsRED (mt-DsRED) and cytoplasmic GFP in layer 2/3 pyramidal neurons at 21 days *in vitro* (DIV). (**b**) High magnification of the boxed dendritic region labeled B in panel A. (**c**) High magnification of the boxed axonal region labeled C in panel A. Notice the striking difference in morphology between elongated, fused mitochondria in dendrites and short mitochondria in the axon. (**d**) Layer 2/3 cortical neurons at (P)ostnatal day 21 were sparsely labeled *in vivo* via *in utero* electroporation with FRT-STOP-FRT plasmids encoding matrix targeted YFP (mt-YFP) and cytoplasmic filler mTAGBFP2. Sparseness was controlled by lowering the concentration of a Flp-e encoding plasmid until single cell resolution was achieved. (**e**) High magnification of the boxed region in panel D. (**f**) The mt-YFP channel of the boxed region of panel D. (**g**) Quantification of mitochondrial length in the axons and dendrites of cortical neurons both *in vivo* and *in vitro*, demonstrating that mitochondria in the axon are much shorter than mitochondria in the dendrites. (**h**) Quantification of the percent of the axonal or dendritic segment occupied by mitochondria both *in vivo* and *in vitro*, demonstrating that mitochondria occupy a much larger percentage of the dendritic arbor than of the axon in cortical neurons. Data is represented by box plots displaying minimum to maximum values, with the box denoting 25^th^, 50^th^ (median) and 75^th^ percentile. n_axons in vivo_ = 21 axons, 235 mitochondria; n_dendrites in vivo_ = 26 dendrites, 267 mitochondria; n_axons in vitro_ = 37 axons, 240 mitochondria; n_dendrites in vitro_ = 43 dendrites, 100 mitochondria. Kruskal-Wallis test with Dunn‘s multiple comparisons test, p<0.0001 for both length and occupancy *in vitro* and *in vivo*.

### Axonal Mitochondria Size is Regulated by Mitochondrial Fission Factor (MFF)

Based on publicly available RNA-seq data from the cerebral cortex and hippocampus ^50,51^ only two of the four known mediators of mitochondrial fission (so called Drp1 ‘receptors’) are highly expressed in excitatory pyramidal neurons: Mitochondrial Fission Factor (*Mff*) and Mitochondrial Fission 1 (*Fis1*). To determine if MFF-and/or FIS1-mediated fission are required for the regulation of mitochondrial size in the axon, we first validated knockdown constructs for both mouse *Mff* and *Fis1* via western blot and immunocytochemistry (**Fig. S1**). Following *ex utero* or *in utero* electroporation of the validated shRNA constructs for *Mff* along with plasmid DNA encoding cytoplasmic filler (tdTomato) and a mitochondria-targeted fluorescent protein (mt-YFP) at E15.5 and visualizatison of neurons at 21DIV or P21, mitochondrial length and occupancy along the axon were dramatically increased compared to control both *in vitro* and *in vivo* (**Fig. 2a-d** and **Fig. 2i-n**). This increase in mitochondrial length and occupancy is completely rescued by the co-expression of a cDNA plasmid encoding human MFF, which is impervious to the shRNA directed against mouse *Mff* (**Fig. 2e-h**), demonstrating that this effect is not the result of an ‘off-target’ effect of the shRNA. In contrast, *Fis1* knockdown had no effect on axonal mitochondrial length but did show a small but significant increase in mitochondrial occupancy which could be consistent with its recently proposed role in mitophagy ^31^ (**Fig. 2o-s**). Interestingly, *Mff* knockdown had no effect on mitochondrial length or occupancy in dendrites *in vivo* suggesting a low level of basal MFF activity in this compartment (**Fig. S2**). These results establish that MFF, but not FIS1, is required for the maintenance of small mitochondrial size in the axon. Interestingly, MFF is present on both somatodendritic and axonal mitochondria (**Fig. S1c-n**) suggesting that fission is regulated in a manner other than localization of MFF to a particular set of mitochondria (See Discussion).

**Figure 2:**
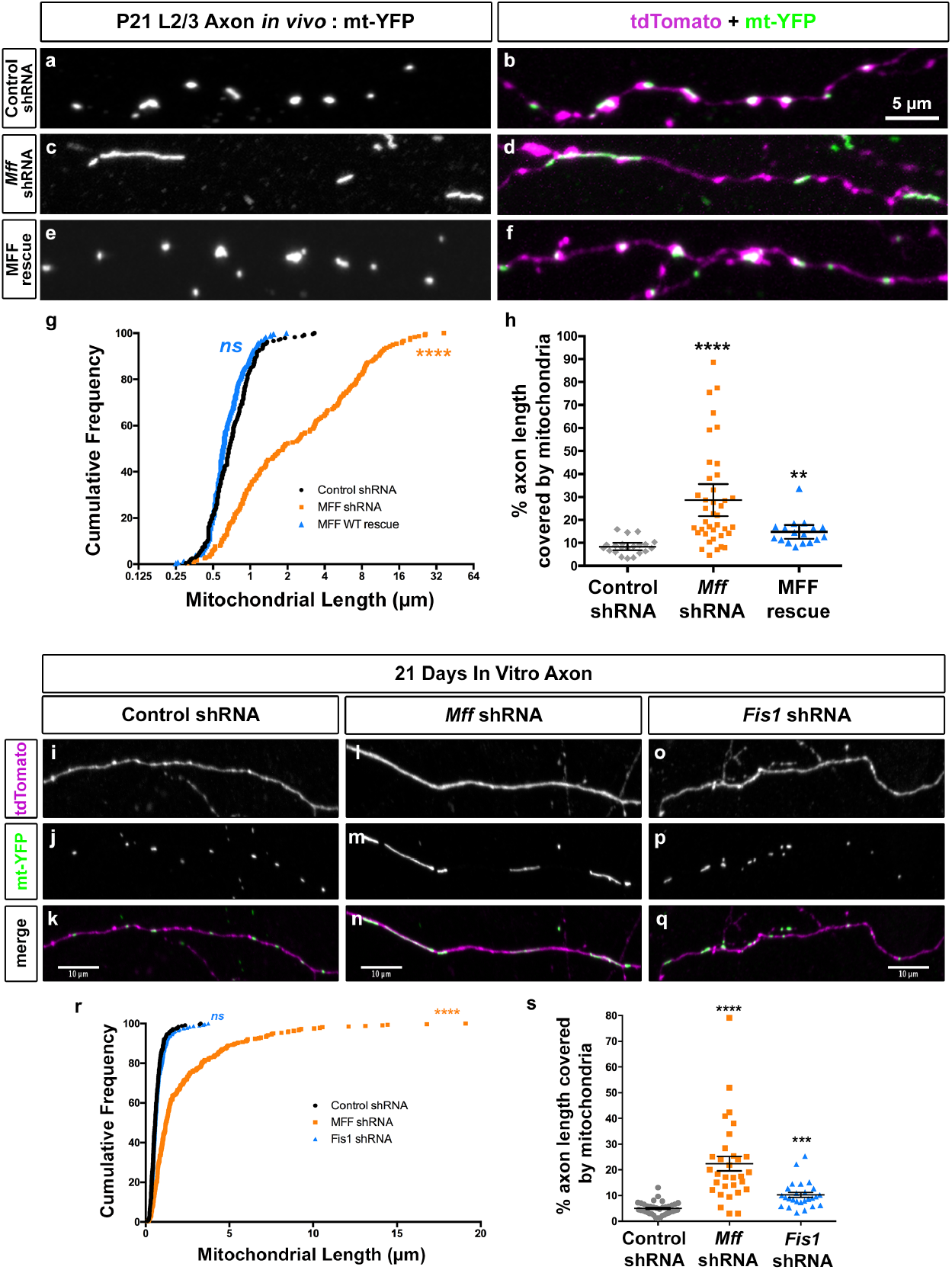
Loss of Mitochondrial Fission Factor (MFF) increases axonal mitochondrial size. (**a**) Mitochondria morphology visualized in a P21 axon after *in utero* electroporation of mt-YFP, cytoplasmic tdTomato, empty pCAG vector, and control shRNA. (**b**) Merge of the mt-YFP and tdTomato channels. (**c**) Mitochondria morphology visualized in a P21 axon after *in utero* electroporation of mt-YFP, cytoplasmic tdTomato, empty pCAG vector, and a mixture of *Mff* shRNAs. (**d**) Merge of the mt-YFP and tdTomato channels. (**e**) Mitochondria morphology visualized in a P21 axon after *in utero* electroporation of mt-YFP, cytoplasmic tdTomato, pCAG Flag-hMFF, and a mixture of *Mff* shRNAs. (**f**) Merge of the mt-YFP and tdTomato channels. (**g**) Cumulative distribution of the number of mitochondria at various sizes under the three conditions above. There is a clear shift to larger mitochondria upon loss of MFF which can be rescued by reintroduction of the shRNA impervious human *MFF*, establishing that MFF is the main regulator of mitochondrial size along the axon. (**h**) Quantification of mitochondrial occupancy of the axon showing that loss of MFF activity leads to an increase of axonal occupancy. Data is represented as a scatter plot with mean ± sem. n_control shRNA_ = 21 axons, 235 mitochondria; n_MFF shRNA_ = 39 axons, 237 mitochondria; n_MFF rescue_= 17 axons, 237 mitochondria. Kruskal-Wallis test with Dunn‘s multiple comparisons test; length: p<0.0001 for control vs. shRNA and rescue vs. shRNA, p= no significance for control vs. rescue; occupancy: p<0.0001 for control vs. shRNA, p<0.01 for control vs. rescue, p= no significance for shRNA vs. rescue. (**i-k**) Representative images of an axon from a neuron electroporated with mt-YFP, tdTomato and control shRNA via *ex utero* electroporation at E15.5, and stained at 21DIV with antibodies for GFP and tdTomato. (**l-n**) Representative images of an axon from a neuron electroporated with mt-YFP, tdTomato and a mixture of *Mff* shRNAs via *ex utero* electroporation at E15.5, and stained at 21DIV with antibodies for GFP and tdTomato. (**o-q**) Representative images of an axon from a neuron electroporated with mt-YFP, tdTomato and a mixture of *Fis1* shRNAs via *ex utero* electroporation at E15.5, and stained at 21DIV with antibodies for GFP and tdTomato. (**r**) Cumulative frequency of mitochondrial length upon *Mff* or *Fis1* shRNA mediated knockdown in 21DIV axons. (**s**) Quantification of percent of axon length occupied by mitochondria. Data is represented as a scatter plot with mean ± sem where each dot represents an axonal segment. Each quantification is from at least 3 independent electroporated mice. n_control shRNA_ = 37 axons, 240 mitochondria; n_MFF shRNA_ = 31 axons, 216 mitochondria, n_Fis1 shRNA_ = 26 axons, 300 mitochondria. Kruskal-Wallis test with Dunn’s multiple comparisons test. length: **** p<0.0001 for control vs. *Mff* shRNA, p= no significance for control vs. *Fis1* shRNA; occupancy: p<0.0001 for control vs. *Mff* shRNA, p<0.01 for control vs. *Fis1* shRNA.

### Reduced MFF Activity Produces Elongated Axonal Mitochondria Via Increased Mitochondrial Length Upon Axonal Entry as well as Decreased Fission Along the Axon Shaft

To better characterize the role of MFF in regulating axonal mitochondrial size, we used the mitochondrial matrix-targeted, photo-convertible fluorescent protein mEos2 ^52,53^ which allowed us to quantify mitochondrial fission and fusion. We employed two distinct imaging strategies to determine where MFF activity is required for axonal mitochondrial size maintenance; (1) photo-conversion of mitochondria located in the cell body followed by time-lapse imaging of mitochondrial entry into the axon, or (2) photo-conversion of a small fraction of mitochondria located along the axon shaft followed by time-lapse imaging and probing of fusion and fission events (**Fig. 3a, Fig. S3a**). Time-lapse imaging of mitochondrial entry into the axon following photo-conversion in the cell body revealed that loss of MFF affects both the frequency of axonal entry as well as their size. On average ~8 mitochondria (8.1 ± 1.7) entered the axon from the cell body per hour in control neurons, but upon *Mff* knockdown this decreased by more than half (2.8 ± 0.78) (**Fig. S3b-d**). Concurrently, the length of mitochondria entering the axon increased four-fold upon *Mff* knockdown compared to control neurons (**Fig. S3e, Movie S1**), establishing that MFF activity in the cell body is required for proper axonal mitochondria size.

**Figure 3:**
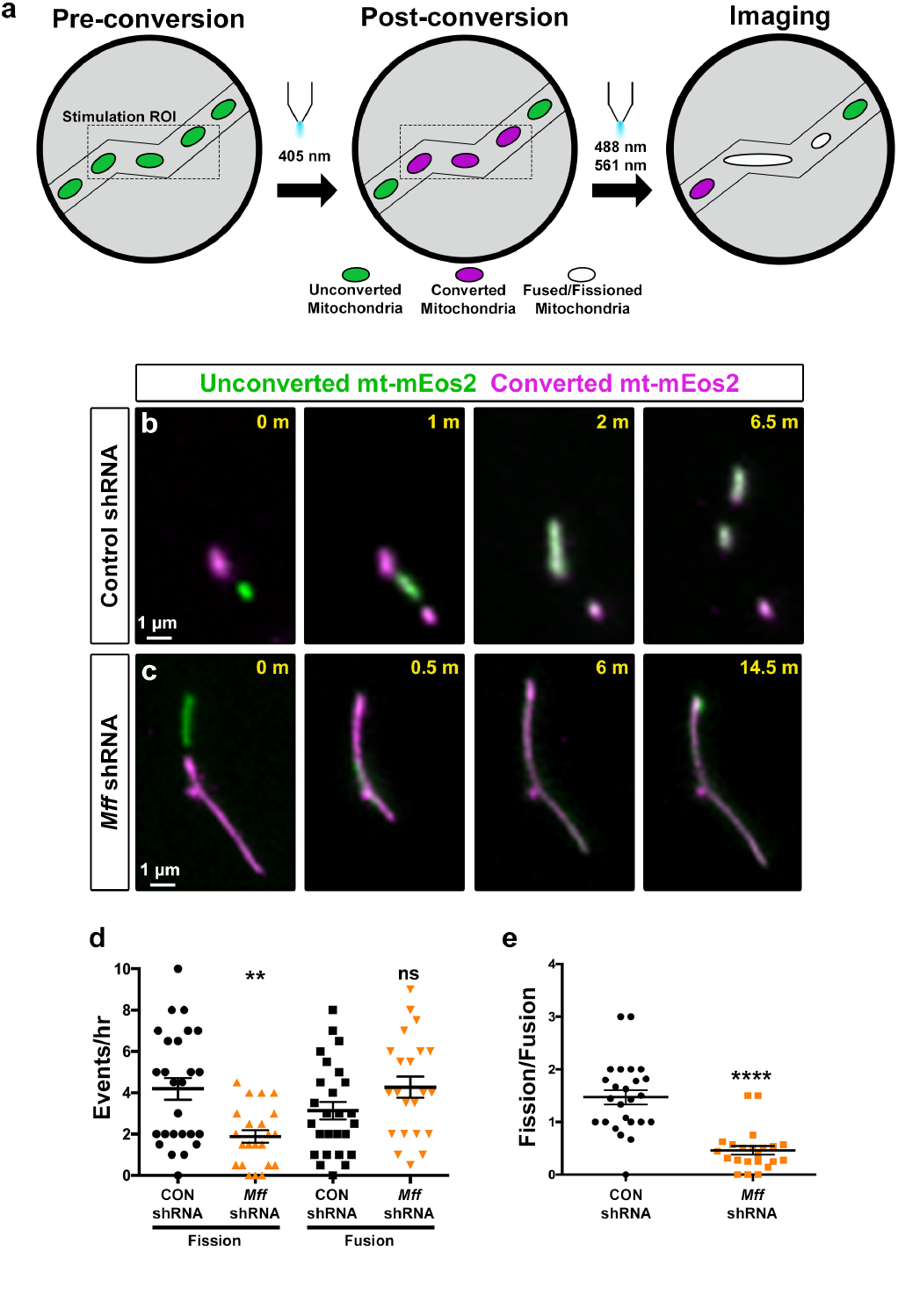
Loss of MFF reduces axonal fission and reverses fission/fusion balance. (**a**) Schematic of the imaging paradigm used for measuring axonal fission and fusion via mitochondria-targeted mEos2 (mt-mEos2), a photoconvertible fluorescent probe. (**b**) Selected timeframes of a mitochondrial fusion and fission event along the axon of a 7DIV neuron *ex utero* electroporated with control shRNA and mt-mEos2. (**c**) Selected timeframes of a mitochondrial fusion event along the axon of a 7DIV neuron *ex utero* electroporated with *Mff* shRNA and mt-mEos2. Fusion events are most frequently coupled with a subsequent fission event in control axons, whereas in MFF-deficient axons, fission is significantly less frequent following a fusion event. See also Movie S1. (**d**) Quantification of the frequency of fission and fusion (#events per hour) per axon segment. Data is represented as a scatter plot with mean ± sem. p=0.0019 for control vs. *Mff* shRNA fission, p=0.1135 for control vs. *Mff* shRNA fusion. (**e**) Quantification of the fission to fusion ratio per axon segment. Data is represented as a scatter plot with mean ± sem. Pπ.0001 for control vs. *Mff* shRNA fission/fusion ratio. Mann-Whitney test. n_control shRNA_ = 26 axons; n_MFF shRNA_ = 22 axons. In both D and E, each dot is a single axon segment. Quantification done from at least 3 independent cultures in each condition

To determine if MFF-dependent fission is also required along the axon for the maintenance of mitochondrial size, we performed time-lapse imaging following photo-conversion of a small subset of mitochondria along the axon shaft. In control axons, the hourly rate of fission (4.2 ± 0.53) and fusion (3.1 ± 0.42) slightly favors fission. However, upon loss of MFF activity, axonal fission is much less likely (1.8 ± 0.30) while fusion is slightly more likely (4.2 ± 0.52) (**Fig. 3b-d**) leading to a dramatic reversal in the fission/fusion ratio (1.5 013 (control shRNA) vs. 0.46 ± 0.08 (*Mff* shRNA) **Fig. 3d, Movie S2**). These results demonstrate that MFF is required both for regulation of mitochondrial size during axonal entry as well as along the axon shaft for maintenance of small mitochondrial size.

### *Mff* Knockdown Disrupts Terminal Axon Branching

To determine the consequence of decreased MFF expression on axonal development, we performed unilateral cortical *in utero* electroporation (IUE) of the validated *Mff* shRNA constructs along with a plasmid encoding cytoplasmic tdTomato at E15.5, and collected coronal sections at P21 to visualize axonal growth and branching. In control brains, axons grew across the corpus callosum reaching the contralateral hemisphere where they invaded the upper layers forming a stereotyped branching pattern in layers 2/3 and 5 (**Fig. 4a-c**) ^13,48,49^. Knockdown of *Mff* did not affect neuronal migration, axon formation and axon growth across the midline. Axons from *Mff* knockdown neurons reached their target territory on the contralateral cortex all the way to superficial layers 2/3. However, terminal axon branching was dramatically reduced, especially in layers 2/3 (**Fig. 4d-f**). This striking reduction of cortical axon branching was rescued upon reintroduction of shRNA-impervious human *MFF* validating the specificity of our shRNA directed against *Mff* (**Fig. 4g-k**). These results emphasize the importance of MFF-dependent fission for the control of axonal mitochondria size and axonal branching *in vivo*.

**Figure 4:**
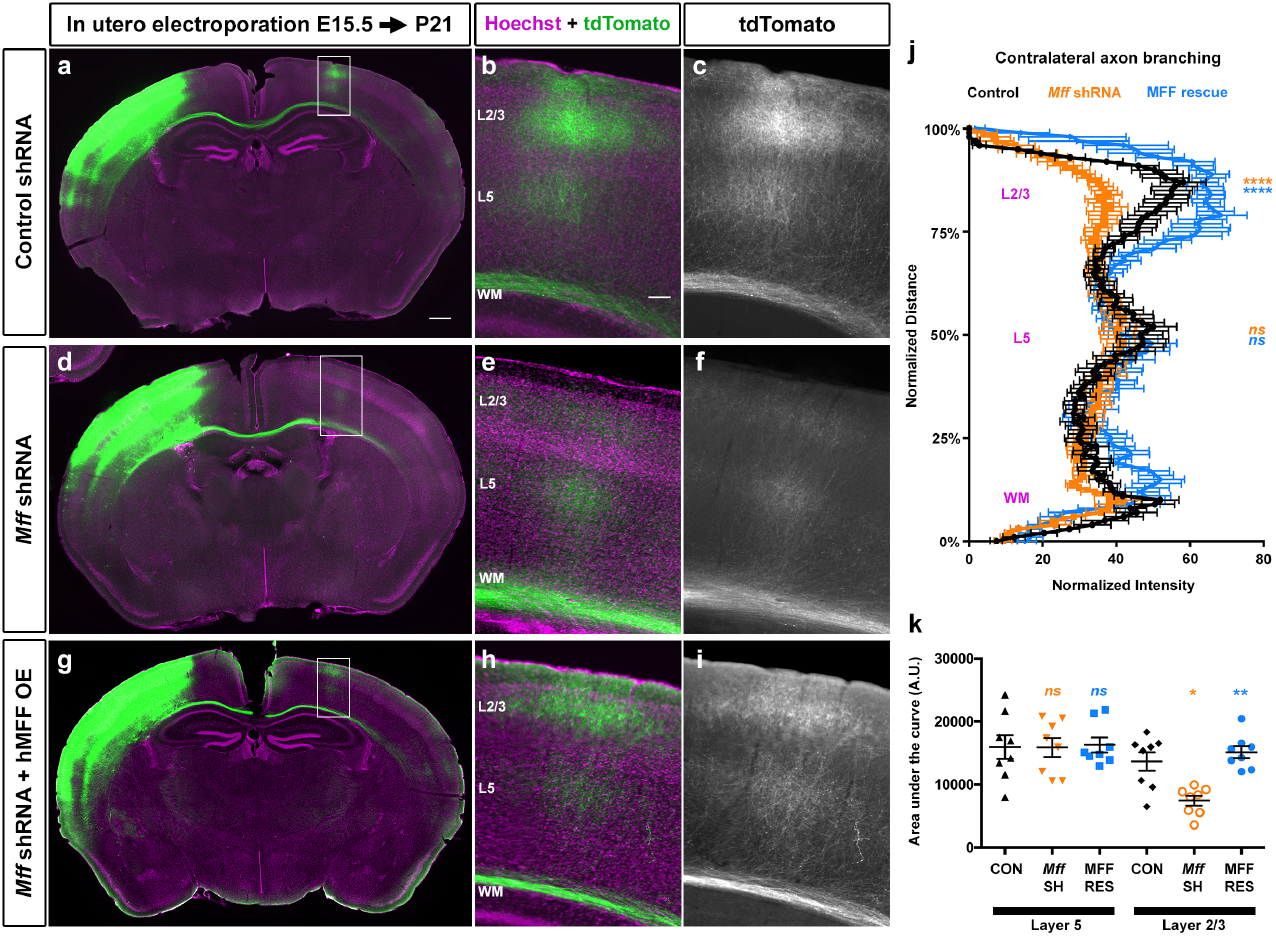
Loss of MFF dramatically reduces contralateral axon branching. (**a**) Low magnification of a P21 mouse brain (coronal section) following cortical *in utero* electroporation with tdTomato, empty pCAG vector and control shRNA. (**b**) Higher magnification of the box in panel A. (**c**) The tdTomato channel only from panel B showing the axon branching pattern of contra-laterally layer 2/3 cortical axons which branch densely in layer 2/3 and layer 5 but avoid branching in layer 4 and 6. (**d**) Low magnification of a P21 coronal section following *in utero* electroporation with tdTomato, empty pCAG vector and a mixture of *Mff* shRNAs (TRC539 and 665; see **Fig. S1A** for validation). (**e**) Higher magnification of the box in panel A. (**f**) The tdTomato channel only from panel E. (**g**) Low magnification of a P21 coronal section following cortical *in utero* electroporation with tdTomato, pCAG::HA-hMFF (rescue construct) and a mixture of *Mff* shRNAs. (**h**) Higher magnification of the box in panel A. (**i**) The tdTomato channel only from panel H. (**j**) Quantification of tdTomato fluorescence along the radial axis of the contralateral cortex (mean ± sem, two-way analysis of variance (ANOVA)) confirming decreased contralateral terminal branching upon MFF loss, and rescue upon reintroduction of shRNA impervious hMFF. ****p<0.0001 and n.s., not statistically significant. (**k**) Quantification of the area under the curves shown in J. Data is represented as a scatter plot with mean ± sem. N_control shRNA_ = 4 brains; n_MFF shRNA_ = 4 brains; n_MFF rescue_ = 4 brains. Each dot represents a single section analyzed. Kruskal-Wallis test with Dunn’s multiple comparisons test. n.s., p003E0.05; * p<0.05;** p<0.01 and **** p<0.0001. In panels J and K, orange characters are for statistical comparisons between control and *Mff* shRNA; blue characters are for statistical comparisons between control and *Mff* shRNA+h*MFF* cDNA rescue condition.

### Reduced MFF Activity Does Not Affect Presynaptic Mitochondrial Capture

Based on our previous work demonstrating the importance of presynaptic mitochondrial capture for terminal axonal branching ^13^, we tested whether altered mitochondrial size had any effect on axonal mitochondrial transport and/or their capture at presynaptic sites. As previously shown mitochondrial motility *in vitro* and *in vivo* progressively decreases along the axons of pyramidal neurons *in vitro* and *in vivo* ^53–55^, so we quantified mitochondrial motility at both 7DIV and 21DIV (**Fig. S4a-b**, **Movie S3**). We observed a modest but significant increase in the number of stationary mitochondria at 7DIV in MFF-deficient axons compared to control. Strikingly as seen in Movie S3, these elongated axonal mitochondria are clearly capable of sustained trafficking along the axon (**Fig. S4c-d**). At 21DIV, the vast majority of these elongated mitochondria become immobilized at specific points along the axon as observed in control axons (94.8 ± 3.53% versus 91.3 ± 5% in control). To determine if altering mitochondrial size along the axon affected their capture at presynaptic boutons *in vivo*, we performed unilateral cortical *in utero* electroporation of the validated shRNA constructs for *Mff* along with plasmid DNA encoding cytoplasmic tdTomato (cell filler), mt-YFP (mitochondrial marker) and VGLUT1-HA (presynaptic marker) at E15.5 and collected coronal sections at P21 to visualize co-localization of mitochondria and VGLUT1 positive presynaptic sites. Interestingly, elongation of axonal mitochondrial size had no significant effect on localization to presynaptic sites or the density of VGLUT1 presynaptic boutons along layer 2/3 cortical axons (**Fig. S4e-h**). These results demonstrate that elongation of axonal mitochondria size does not impact their capture at presynaptic boutons and suggested that we should instead focus on potential changes in mitochondrial function that might accompany the observed increase in mitochondrial size upon knockdown of *Mff*.

### Reduced MFF Activity and Elongation of Axonal Mitochondria Does Not Decrease Mitochondrial Membrane Potential, Mitochondrial ATP Levels or Redox Potential

We first tested if either mitochondrial membrane potential or mitochondrial ATP levels were affected following a reduction in MFF levels and increased mitochondrial size along cortical axons. To determine if these elongated axonal mitochondria maintain their membrane potential, we used Tetramethylrhodamine, Methyl Ester (TMRM, 20nM). We electroporated cortical neurons with either control shRNA or *Mff* shRNA and unique combinations of mitochondrial and cytoplasmic fluorescent proteins (mt-BFP and Venus for control vs. mt-YFP and mTAGBFP2 for *Mff* knockdown), and mixed control and knockdown neurons at a 1:1 ratio (**Fig. S5**). This approach allowed us to image mitochondrial membrane potential in control and *Mff* knockdown neurons under the exact same culture conditions. Upon quantification of the Fm/Fc ratio for individual mitochondria along the axon ^56^, the longer axonal mitochondria upon *Mff* knockdown actually have an increased ratio (data not shown) but after accounting for mitochondrial area there is no change in the Fm/Fc ratio versus the control mitochondria (15.0 ± 2.63 in *Mff* knockown vs 13.8 ± 1.08 for control, **Fig. 5a-b**).

**Figure 5:**
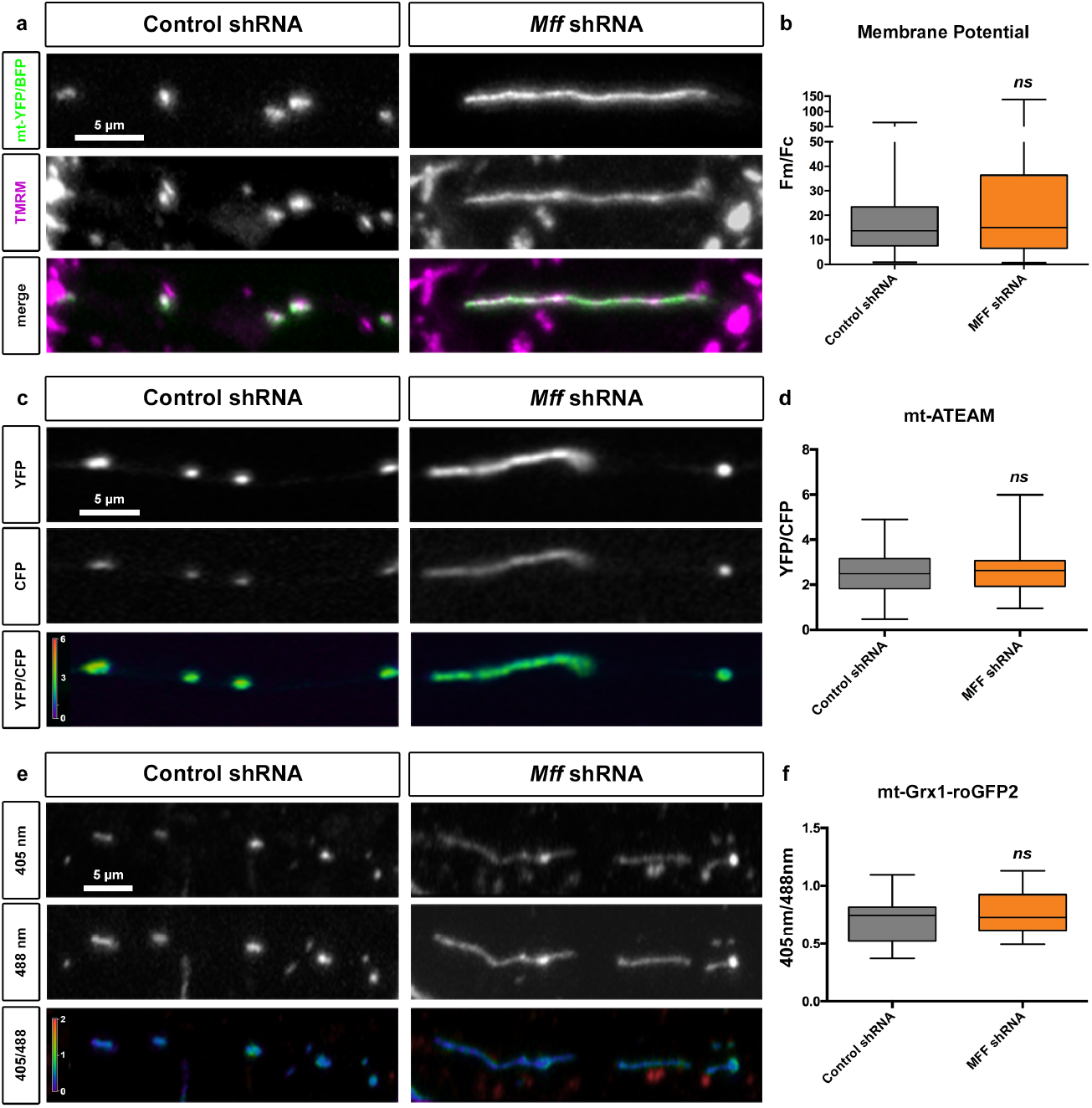
Loss of MFF does not alter mitochondrial membrane potential, mitochondrial ATP levels or redox potential of mitochondrial matrix. (**a**) Representative images from axons of neurons electroporated with mt-mTAGBFP2, Venus (cell filler) and control shRNA (left panels) or mt-YFP, mTAGBFP2 (cell filler) and *Mff* shRNA mixture (right panels) via *ex utero* electroporation at E15.5, and treated with 20nM TMRM at 7DIV. (**b**) Quantification of((F_mitochondria_/F_cytoplasm_)/(Mitochondrial Area)) for TMRM at 7DIV. Data is represented at minimum to maximum box plots, with the box denoting 25^th^, 50^th^ (median) and 75^th^ percentile. n_control shRNA_ = 7 axons, 125 mitochondria; n_MFF shRNA_ = 11 axons, 104 mitochondria, n.s. p>0.05 according to non-parametric Mann-Whitney test. (**c**) Representative images from axons of neurons electroporated with mt-ATEAM, mCardinal and control shRNA (left panels) or mt-ATEAM, mScarlet and *Mff* shRNA mixture (right panel) via *ex utero* electroporation at E15.5 and visualized at 21DIV. (**d**) Quantification of (F_YFP_/F_CFP_) for ATEAM at 21DIV. Data is represented at minimum to maximum box plots, with the box denoting 25^th^, 50^th^ and 75^th^ percentile. n_control_ shRNA = 13 axons, 176 mitochondria; n_MFF shRNA_ = 18 axons, 145 mitochondria, p= 0.4504 by Mann-Whitney. (**e**) Representative images from axons of neurons electroporated with mt-Grx1-roGFP2 and control shRNA (left panels) or mt-Grx1-roGFP2 and *Mff* shRNA mixture (right panels) via cortical *ex utero* electroporation at E15.5 and visualized at 21DIV. (**f**) Quantification of (F_405nm_/F_488nm_) for roGFP2 at 21DIV. Data is represented at minimum to maximum box plots, with the box denoting 25^th^, 50^th^ and 75^th^ percentile. n_control_ shRNA = 28 axons, 405 mitochondria; n_MFF shRNA_ = 22 axons, 238 mitochondria, p>0.05 according to Mann-Whitney test.

Taking a similar approach, we measured intra-mitochondrial ATP levels using a genetically-encoded probe, ATEAM1.03, targeted to the mitochondrial matrix ^57,58^ (mt-ATEAM and mCardinal for control vs. mt-ATEAM and mScarlet for *Mff* knockdown). Again, we found no significant change in the YFP/CFP ratio for mitochondria upon *Mff* knockdown (2.55 ± 0.07) versus control (2.38 ± 0.08) imaged with the same settings and culture conditions at 21DIV (**Fig. 5c-d**).

Finally, we tested if there was a change in the redox potential of elongated, Mff-deficient mitochondria using a genetically-encoded redox-sensitive probe targeted to the mitochondrial matrix (mt-roGFP2; ^59,60^). Upon *Mff* knockdown, there was no significant change in the ratio of 405nm/488nm excitation (0.77 ± 0.04) compared to neurons expressing control shRNA (0.70 ± 0.04) at 21DIV (**Fig. 5e-f**). These results strongly argue that reducing MFF-dependent mitochondrial fission in axons does not affect their ability to perform oxidative phosphorylation and therefore their ability to generate ATP or their redox state.

### Axonal Mitochondrial Elongation upon *Mff* Knockdown Increases Mitochondrial Calcium Uptake at Presynaptic Sites

Based on previous results from our lab and others demonstrating that mitochondrial Ca^2+^ uptake plays a critical role in regulating presynaptic release at *en passant* boutons of pyramidal neurons ^15,16^, we tested if mitochondrial Ca^2+^ uptake was affected upon *Mff* knockdown. Our hypothesis was that the total amount of Ca^2+^ uptake capacity of mitochondria is in part dependent on their total matrix volume. To monitor presynaptic mitochondrial Ca^2+^ dynamics ([Ca^2+^]_m_), we used a mitochondrial matrix targeted genetically-encoded Ca^2+^ sensor, mt-GCaMP5G ^15,61^, and introduced this with VGLUT1-mCherry and mt-mTagBFP2 to cortical neurons using *ex utero* electroporation at E15.5. At 17-23DIV, we imaged intra-mitochondrial Ca^2+^ dynamics before and following stimulation of presynaptic release with 20 action potentials (AP) at 10Hz using a concentric bipolar electrode ^15,62^. During stimulation, Ca^2+^ import occurs at points of these long mitochondria in direct contact with presynaptic sites, followed by diffusion of mitochondrial Ca^2+^ along the mitochondria in *Mff* knockdown neurons (**Fig. 6b, Fig. S6 and Movie S4**). In addition, long mitochondria show significantly faster extrusion of mitochondrial Ca^2+^ after stimulation compared to control mitochondria (t=13.2 ± 1.5s of versus 8.2 ± 1.1s in control, **Fig. 6a-b and e**). To determine if these elongated mitochondria accumulate more Ca^2+^, we measured the integrated intensity of mt-GCaMP5G over the entire length of individual mitochondria associated with single presynaptic VGLUT1+ boutons. Interestingly, elongated presynaptic mitochondria import significantly higher amounts of Ca^2+^ than punctate mitochondria (as measured by area under the curve or total charge transfer: 2.3×10^5^ ± 0.41×10^5^ versus 1.4×10^5^ ± 0.19×10^5^ in control, **Fig. 6c-d**).

**Figure 6.**
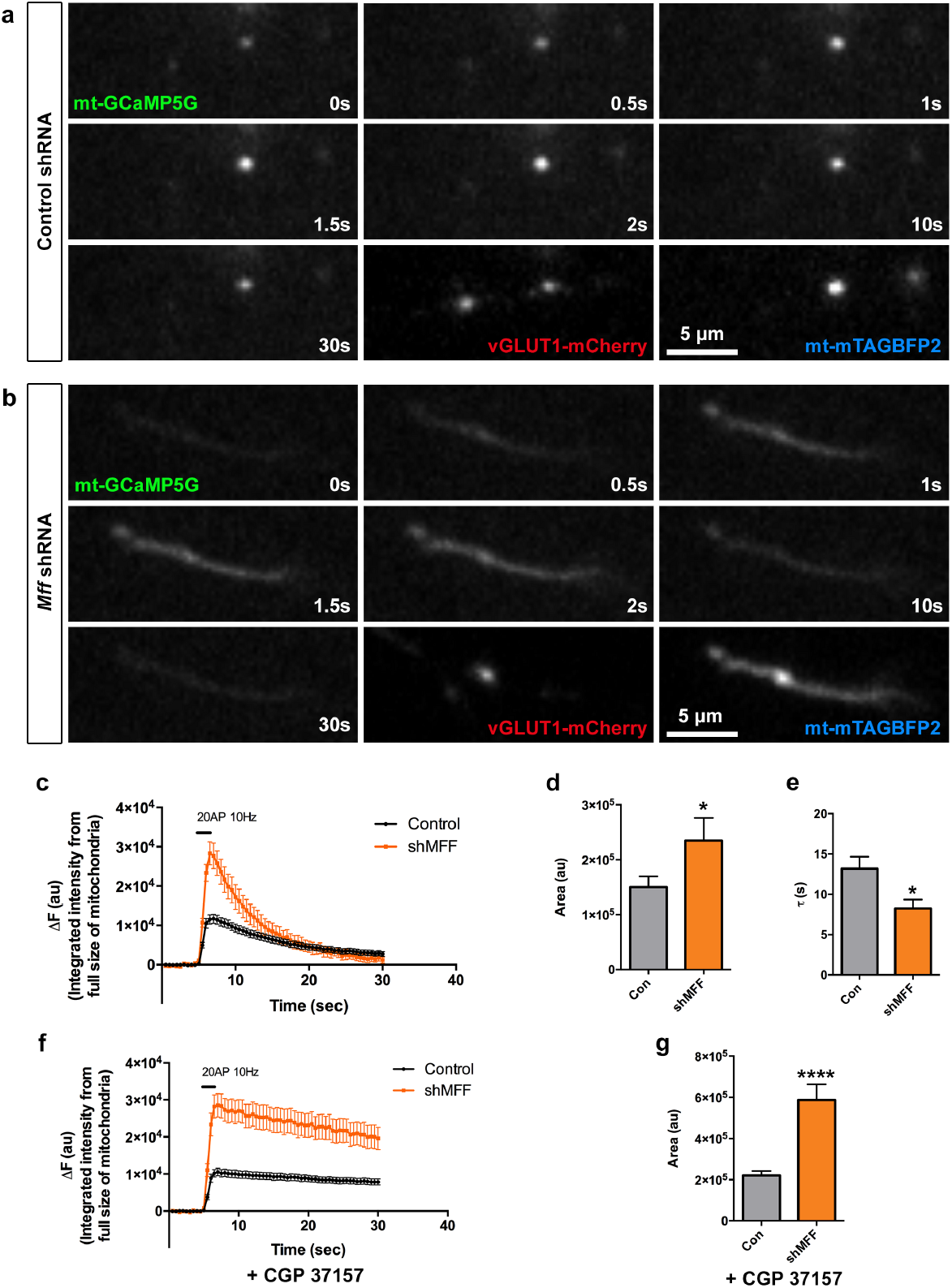
Elongated mitochondria in *Mff* knockdown axons uptake more Ca^2+^ upon evoked neurotransmitter release from presynaptic sites. (**a-g**) Presynaptic mitochondrial Ca^2+^ was monitored using mitochondria-targeted GCaMP5G (mt-GCaMP5G) with VGLUT1-mCherry and mt-mTagBFP2 in cortical cultured neurons following *ex utero* electroporation at E15.5 and imaged at 17-23DIV. (**a-b**) Cropped time lapse images of mt-GCaMP5G with repetitive stimulation (20AP at 10Hz) in control and *Mff* knockdown axons. Mitochondrial Ca^2+^ diffuses along elongated mitochondria from a presynaptic site in an *Mff* knockdown axon (See Figure S5 and Movie S4) and extrudes faster than small mitochondria. (**c**) Integrated intensity of mt-GCaMP5G signals from full-length mitochondria associated with single presynaptic sites is plotted with mean ± sem. (**d**) Quantification of [Ca^2+^]_mt_ (area under the curve) show long mitochondria with a single presynaptic bouton in *Mff* knockdown axons accumulate significantly more [Ca^2+^]_mt_ than small mitochondria in control axons. * p<0.05, Mann-Whitney test. (**e**) Long mitochondria show faster decay than control mitochondria. * p<0.05, Mann-Whitney test. (**f-g**) Blocking extrusion of mitochondrial Ca^2+^ by inhibition of mitochondrial Na^+^/Ca^2+^ exchanger (NCLX) causes more accumulation of [Ca^2+^]_mt_ in long mitochondria. Images were captured after CGP 37157 incubation (10μM, 3min) from the same axons in (C). All bar graphs are represented with mean ± sem. n_control_= 7 dishes, 25 mitochondria; n_MFF shRNA_ = 11 dishes, 14 mitochondria. ****p<0.0001, Unpaired t-test.

We next tested if the increased mitochondrial Ca^2+^ uptake we observed in elongated mitochondria upon *Mff* knockdown was impacted by changes in mitochondrial Ca^2+^ extrusion by applying a mitochondrial Na^+^/Ca^2+^ exchanger (NCLX) antagonist, CGP 37157, as NCLX is one of the main mitochondrial Ca^2+^ extrusion mechanisms. Interestingly, this inhibitor effectively delays Ca^2+^ extrusion from mitochondria and actually increases the difference of total Ca^2+^ charge transfer between *Mff* knockdown and control (5.9×10^5^ ± 7.7×10^4^ versus 2.2×10^5^ ± 2.1×10^4^ in control, **Fig. 6f-g**). These results demonstrate that the longer presynaptic mitochondria induced by *Mff* knockdown underlies a significant increase in their Ca^2+^ buffering capacity.

### Increased Uptake of Ca^2+^ by Elongated Mitochondria in *Mff* Knockdown Neurons Decreases Presynaptic Ca^2+^ Levels

To understand the impact of this increased mitochondrial Ca^2+^ uptake, we tested if presynaptic cytoplasmic Ca^2+^ ([Ca^2+^]_cyto_) is affected by the increased Ca^2+^ uptake from mitochondria in *Mff* knockdown axons. Combining GCaMP5G targeted to presynaptic boutons (VGLUT1-GCaMP5G) with mt-mTagBFP2 and VGLUT1-mCherry allowed us to measure presynaptic Ca^2+^ dynamics at presynaptic boutons associated with mitochondria^15^. All three plasmids with either control or *Mff* shRNA were introduced by *ex utero* cortical electroporation at E15.5 and imaged in axons of pyramidal neurons at 17-23DIV using the stimulation protocol described above (20APs at 10Hz). Interestingly, the peak of [Ca^2+^]_cyto_ signals from long mitochondria-associated presynaptic boutons was 20% lower (0.46 ± 0.025 versus 0.56 ± 0.02 in control, **Fig. 7a-b**). In addition, both the peak values of presynaptic [Ca^2+^]_cyto_ and the total charge transfer (area under the curve) were significantly reduced in *Mff* knockdown neurons (**Fig. 7c-d**). These results show that the increased [Ca^2+^]_m_ characterizing MFF-deficient mitochondria significantly impacts presynaptic [Ca^2+^]_cyto_ clearance.

**Figure 7.**
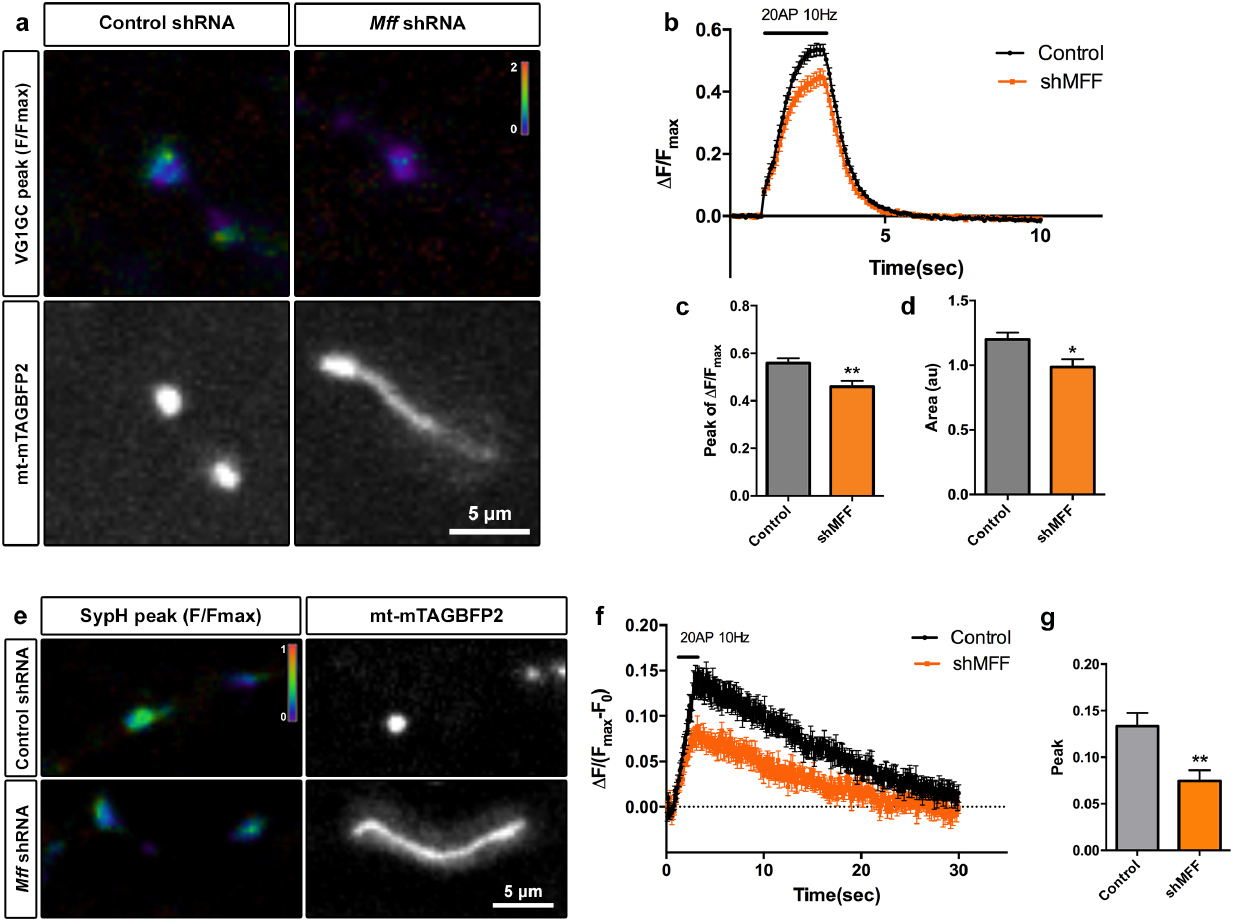
Presynaptic boutons associated with long mitochondria in MFF-deficient axons show decreased Ca^2+^ accumulation and reduced evoked neurotransmitter release. (**a-d**) Presynaptic Ca^2+^ dynamics in *Mff* knockdown cortical neurons were monitored using VGLUT1-GCaMP5G with mt-mTagBFP2 and VGLUT1-mCherry at 17-23DIV. (**a**) Representative images show VGLUT1-GCaMP5G peak at 20APs (10Hz) and mt-mTagBFP2. VGLUT1-GCaMP5G is displayed by ratio view normalized (ΔF/F_max_) by F_max_ values obtained following ionomycin (5µM) treatment at the end of each imaging session. (**b-d**) Presynaptic boutons from MFF-deficient neurons have significantly decreased peak value and total charge transfer (area under the curve). All graphs are represented with mean ± sem. n_control_= 9 dishes, 38 boutons; n_MFF shRNA_ = 11 dishes, 24 boutons. p=0.001 for peak, p=0.018 for area, Mann-Whitney test. (**e-g**) Presynaptic release properties linked to long mitochondria were monitored using synaptophysin-pHluorin and mt-mTagBFP2 in cultured neurons at 20-23DIV. (**e**) Representative images of sypH peak normalized by F_max_ obtained during NH_4_Cl (50mM) incubation. (**f-g**) Presynaptic release associated with long mitochondria in *Mff* knockdown axons is significantly reduced during 20APs (10Hz) compared to control (F). Quantification of SypH peak normalize values (ΔF/(F_max_-F_0_)) show significantly reduced neurotransmitter vesicle exocytosis during stimulation of presynaptic release with 20AP@10Hz. All graphs are represented with mean ± sem. n_control_ = 16 dishes, 47 boutons; n_MFF shRNA_ = 24 dishes, 33 boutons. p=0.0037, Unpaired t-test.

### Presynaptic Release at Sites of Elongated Mitochondria is Impaired in *Mff* Knockdown Neurons

Presynaptic Ca^2+^ influx through voltage-gated Ca^2+^ channels (VGCC) triggers synaptic vesicle exocytosis, therefore the amount of presynaptic Ca^2+^ quantitatively regulates neurotransmitter release at individual boutons ^15,63^. To monitor presynaptic release at individual boutons, we employed pHluorin-tagged synaptophysin (syp-pHluorin) ^64^. In short, pH-sensitive GFP, pHluorin, is fused to the luminal domain of synaptophysin, and syp-pHluorin fluorescence is quenched because of the low/acidic luminal pH of neurotransmitter vesicles ^65^. When exposed to extracellular pH following vesicle exocytosis, the luminal pH equilibrates closer to pH 7 and pHluorin emits fluorescence. We introduced the syp-pHluorin, synaptophysin-mCherry (constitutive presynaptic marker), and mt-mTagBFP2 with either control or *Mff* shRNA using *ex utero* electroporation at E15.5 and imaged at 20-23DIV. Consistent with decreased presynaptic [Ca^2+^]_cyto_ levels, presynaptic boutons associated with long mitochondria in *Mff* knockdown axons displayed significantly decreased evoked neurotransmitter vesicle release (0.074 ± 0.012 versus 0.13 ± 0.014 in control, **Fig. 7e-g**). These results demonstrate that, in cortical axons, regulation of mitochondrial size by MFF-dependent fission is critical for proper presynaptic Ca^2+^ homeostasis, neurotransmitter release and results in decreased terminal axon branching *in vivo*.

## Discussion

In the present study, we uncovered a novel mechanism by which neurons regulate presynaptic release and axonal development. Our results demonstrate that loss of MFF activity increased mitochondria length by reducing fission for both mitochondria entering the axon as well as along the axonal shaft. Strikingly, mitochondrial elongation did not affect mitochondrial trafficking, presynaptic positioning, membrane potential, or their ability to produce ATP; however, it dramatically increased their Ca^2+^ buffering capacity. This size-dependent increase in mitochondrial Ca^2+^ uptake lessened presynaptic cytoplasmic Ca^2+^ levels causing reduced neurotransmitter release upon evoked activity and resulted in decreased terminal axon branching *in vivo* (**Fig. S7**).

These results suggest a two-step model for MFF-dependent regulation of axonal mitochondrial size where MFF activity is required both at the time of axonal entry as well as for maintenance along the length of the axon. Our data reveals that in control cortical axons, mitochondria enter with an average size of ~1µm, and maintain this short length along the axon by coupling almost every fusion event to a fission event. Our results represent the first evidence that axonal mitochondrial size is regulated before or upon entry into the axon, and raises intriguing possibilities regarding the underlying mechanism(s). Two non-mutually exclusive scenarios exist: (1) mitochondria are “tagged” as axonal (vs. dendritic) mitochondria soon after biogenesis in the cell body; and/or (2) mitochondria allowed to enter the axon are ‘filtered’ based upon size. The most compelling evidence supporting scenario one may be that in hippocampal and cortical neurons axonal and dendritic mitochondria interact with the distinct motor adaptor proteins TRAK1 or TRAK2 respectively ^66,67^. However, the mechanism and site of first interaction between TRAK1/2 and mitochondria remain unknown i.e. do they interact in the cell body to determine specificity or are TRAK1/2 themselves targeted to the axonal or dendritic compartments and engaged locally? Evidence for scenario two and the existence of a size filter only allowing mitochondria of a short length to enter the axon is currently unexplored. However, recent work demonstrating the presence of an actin based filter at the axon initial segment (AIS) ^68,69^ and evidence of myosin motors opposing kinesin-directed mitochondrial movement ^70^ could represent such a mitochondria-size selection filter.

While not the focus of this manuscript, the question of how MFF-dependent fission is regulated differentially in the axon remains poorly understood and should be the focus of further investigations. In fact, it is further complicated by the finding that MFF is present on the outer membrane of both axonal and dendritic mitochondria (**Fig. S1**). Coupled with the results showing no significant change in dendritic mitochondria length upon *Mff* knockdown (**Fig. S2**), this implies that MFF recruitment, activation or interaction with Drp1 must be differentially regulated in the axonal and somatodendritic compartments. Cortical pyramidal neurons could accomplish this via: (1) subcellular targeting of different isoforms of MFF which can be generated through alternative splicing and have different affinities for Drp1 ^32^, (2) MFF can be post-translationally modified and ‘activated’ to increase its ability to recruit Drp1 ^71^, (3) although MiD51 and MiD49 are expressed at extremely low levels in neurons, they may potentially alter the MFF/Drp1 interaction from promoting fission to inhibiting fission ^72,73^, and finally (4) axonal or dendritic Drp1 may be post-translationally modified to increase or reduce its ability to interact with MFF ^10,74–76^.

Our data illustrates that the long mitochondria produced upon *Mff* knockdown possessed a comparable membrane potential and maintained similar levels of matrix ATP (**Fig. 5**); in fact, based on their increased size, we might expect that they actually have an increased capacity to produce ATP. A previous report observed a requirement for activity-driven ATP synthesis in synaptic function ^14^; however, ATP production could only be seen upon a prolonged stimulation condition (600APs at 10Hz). It is now clear that mitochondrial Ca^2+^ buffering is important for synaptic function in more physiological conditions ^15,16^ and we observed significant increases in presynaptic mitochondrial Ca^2+^ uptake upon Mff-knockdown leading to reduced synaptic vesicle release with physiological stimulation regimes (20APs @ 10Hz). In addition, blocking glycolysis or ATP production mainly affected endocytosis of synaptic vesicles (SVs), but not exocytosis ^14^. Layer 2/3 cortical neurons are well known to have less spontaneous activity than other pyramidal neurons ^77,78^; therefore, our results in *Mff*-deficient axons is more likely to be a result of abnormal SV release due to increased mitochondrial Ca^2+^ uptake than altered ATP production.

A recent study showed that the Bcl-x_L_-Drp1 complex regulates SV endocytosis, and MFF may form a complex with these proteins ^79^. Consistent with our data, *Mff* knockdown in hippocampal neurons showed reduced exocytosis of SVs. While the authors of this study normalized the pHluorin signal with F_0_ value from base line signals, we used the F_max_ value obtained by NH_4_Cl incubation in order to normalize for any potential changes in total vesicle pool size. Even with this more optimal normalization method, MFF-deficient neurons still have reduced SV release. Future experiments will be required to determine the importance of both MFF-mediated decreases on synaptic vesicle recycling and increased mitochondrial Ca^2+^ uptake on neurotransmitter release.

In addition to mitochondria, axonal endoplasmic reticulum (ER) was recently shown to be involved in presynaptic Ca^2+^ clearance upon stimulation of neurotransmitter release ^80^. Although inhibition of ER Ca^2+^ uptake can change plasma membrane Ca^2+^ entry through STIM1, a recent 3D-EM study revealed the presence of ER and mitochondria contacts along the axon and specifically at presynaptic sites ^81^. Future investigation is needed to characterize the relative contribution of axonal ER and mitochondria, and ER-mitochondria coupling for presynaptic Ca^2+^ homeostasis.

The role of neuronal activity in axonal development and branching has been well established over the last twenty years. Early work demonstrated that developing neurons require spontaneous activity and spontaneous neurotransmitter release for axonal development ^82–86^ while more recent work has revealed the requirement for activity on both the presynaptic and postsynaptic sides ^48,49,87–90^. Finally, axonal branches which make presynaptic boutons and are synchronized with postsynaptic neuronal activity are more likely to be stabilized when compared to axons with desynchronized activity ^91–93^. Based on these previous results and the observation that layer 2/3 terminal branching is the most affected, it seems likely that the loss of terminal axon branching we observe upon *Mff* knockdown is a result of the decreased neurotransmitter release we observed along these axons.

Taken together, our data demonstrate for the first time that maintenance of small mitochondrial size in CNS axons through MFF-dependent fission is critical to limit presynaptic Ca^2+^ dynamics, neurotransmitter release, terminal axonal branching and therefore proper development of circuit connectivity.

## Author Contributions

Conceptualization, T.L.L., R.S., F.P.; Methodology, T.L.L., S.K., F.P.; Investigation, T.L.L., S.K., A.L.; Writing, T.L.L., S.K., F.P.; Funding Acquisition, T.L.L. and F.P.; Resources, R.S.; Supervision, T.L.L., and F.P.

## Acknowledgements

The authors would like to thank members of the F.P. and R.S. labs for helpful suggestions, technical advice, and critical reading of the manuscript. Funding sources include NIH-R01NS089456 (F.P.), NIH-K99NS091526 (T.L.L.), Human Frontiers Science Program Long-term Fellowship (S.K.), and an Award from the Fondation Roger De Spoelberch (F.P.).

## METHODS

### Mice for in utero electroporation

All animals were handled according to protocols approved by the Institutional Animal Care and Use Committee (IACUC) at Columbia University. Time-pregnant CD1 females were purchased from Charles Rivers. Timed-pregnant hybrid F1 females were obtained by mating inbred 129/SvJ females (Charles Rivers), and C57Bl/6J males (Charles Rivers) in house. At the time of *in utero* electroporation (Embryonic Day 15.5), littermates were randomly assigned to experimental groups without regard to their sex.

### Mice for ex utero electroporation and primary culture

All animals were handled according to protocols approved by the Institutional Animal Care and Use Committee (IACUC) at Columbia University. Time-pregnant CD1 females were purchased from Charles Rivers. At the time of ex utero electroporation (E15.5), littermates were randomly assigned to experimental groups without regard to their sex.

### HEK 293T Cells

Human Embryonic Kidney (HEK) 293T cells were purchased from ATCC.

### Plasmids

pCAG mt-ATEAM was made by placing the DNA encoding mt-ATEAM (gift from Hiromi Imamura) 3’to the CAG promoter via PCR and Infusion cloning (Clonetech). pCAG mt-Grx1-roGFP2 was made by placing the DNA encoding mt-Grx1-roGFP2 (Addgene plasmid #64977) 3’ to the CAG promoter as above. pCAG vGLUT1-HA was created by replacing the mCherry in pCAG vGLUT1-mCherry with an HA tag via restriction digest and PCR. pCAG mTAGBFP2 was created by removing the mitochondrial targeting sequence in pCAG mt-mTAGBFP2 via restriction digest and re-ligation. pCAFNF mTAGBFP2 and mt-YFP were made by PCR of the DNA encoding mTAGBFP2 or mt-YFP and placement 3’ to the CAG FNF cassette (Addgene plasmid #13772) via PCR and Infusion cloning (Clonetech). pCAG HA-mMff and pCAG HA-mFis1 were created via PCR of the DNA encoding mouse *Mff* or mouse *Fis1* from a neuronal mouse cDNA library, and sub-cloned 3’ to the CAG promoter and a HA tag.

### In utero electroporation

A mix of endotoxin-free plasmid preparation (2mg/mL total concentration) and 0.5% Fast Green (Sigma) was injected into one lateral hemisphere of E15.5 embryos using a Picospritzer III (Parker). Electroporation (ECM 830, BTX) was performed with gold paddles to target cortical progenitors in E15.5 embryos by placing the anode (positively charged electrode) on the side of DNA injection and the cathode on the other side of the head. Five pulses of 45 V for 50 ms with 500 ms interval were used for electroporation. Animals were sacrificed 21 days after birth (P21) by terminal perfusion of 4% para-formaldehyde (PFA, Electron Microscopy Sciences) followed by overnight postfixation in 4% PFA. For sparse labeling via *in utero* electroporation, Flp dependent plasmids (pCAFNF, Addgene) were used along with 640 pg/µl of pCAG Flp-e (Addgene).

### Ex utero cortical electroporation

A mix of endotoxin-free plasmid preparation (2-5mg/mL) and 0.5% Fast Green (Sigma) mixture was injected using a Picospritzer III (Parker) into the lateral ventricles of isolated head of E15.5 mouse embryo, and electoporated using an electroporator (ECM 830, BTX) with four pulses of 20V for 100ms with a 500ms interval. Following ex utero electroporation we performed dissociated neuronal culture as described below.

### Primary neuronal culture

Embyonic mouse cortices (E15.5) were dissected in Hank’s Balanced Salt Solution (HBSS) supplemented with HEPES (10mM, pH7.4), and incubated in HBSS containing papain (Worthington; 14U/ml) and DNase I (100 μg/ml) for 20min at 37°C. Then, samples were washed with HBSS, and dissociated by pipetting. Cell suspension was plated on poly-D-lysine (1mg/ml, Sigma)-coated glass bottom dishes (MatTek) or coverslips (BD bioscience) in Neurobasal media (Invitrogen) containing B27 (1x), Glutamax (1x), FBS (2.5%) and penicillin/streptomycin (0.5x, all supplements were from Invitrogen). After 5 to 7 days, media was changed with supplemented Neurobasal media without FBS.

### Immunocytochemistry

Primary culture-Cells were fixed for 10 minutes at room temperature in 4% (w/v) paraformaldehyde (PFA, EMS) in PBS (Sigma), then incubated for 30 minutes in 0.1% Triton X-100 (Sigma), 1% BSA (Sigma), 5% Normal Goat Serum (Invitrogen) in PBS to permeabilize and block nonspecific staining, after washing with PBS. Primary and secondary antibodies were diluted in the buffer described above. Primary antibodies were incubated at room temperature for 1 hour and secondary antibodies were incubated for 30 minutes at room temperature. Coverslips were mounted on slides with Fluoromount G (EMS). Primary antibodies used for immunocytochemistry in this study are chicken anti-GFP (5 μg/ml, Aves Lab – recognizes GFP and YFP), mouse anti-HA (1:500, Covance), rabbit anti-RFP (1:1,000, Abcam – recognizes mTagBFP2, DsRED and tdTomato). All secondary antibodies were Alexa-conjugated (Invitrogen) and used at a 1:2000 dilution. Nuclear DNA was stained using Hoechst 33258 (1:10,000, Pierce)

Endogenous MFF staining-Cells were fixed for 10 minutes at room temperature in 4% PFA in PBS, followed directly with −20°C, 100% methanol at −20°C for 6 minutes. After washing with room temperature PBS, cells were incubated for 30 minutes in 1% BSA (Sigma), 5% Normal Goat Serum in PBS to block nonspecific staining. Primary and secondary antibodies were diluted in the buffer above. Primary antibodies were incubated at room temperature for 1 hour and secondary antibodies were incubated for 30 minutes at room temperature. Coverslips were mounted on slides with Fluoromount G (EMS). Primary antibodies used for this section are chicken anti-GFP (5 μg/ml, Aves Lab – recognizes GFP and YFP), rabbit anti-MFF (1:200, Protein Tech). All secondary antibodies were Alexa-conjugated (Invitrogen) and used at a 1:2000 dilution. Brain sections-Post fixed brains were sectioned via vibratome (Leica VT1200) at 100 µm. Floating sections were then incubated for 2 hours in 0.4% Triton X-100, 1% BSA, 5% Normal Goat Serum in PBS to block nonspecific staining. Primary and secondary antibodies were diluted in the buffer described above. Primary and secondary antibodies were incubated at 4°C overnight. Sections were mounted on slides and coversliped with Aqua PolyMount (Polymount Sicences, Inc). Primary and secondary antibodies are the same as above.

### Imaging

Fixed samples were imaged on a Nikon Ti-E microscope with an A1 confocal. All equipment and solid state lasers (Coherent, 405nm, 488nm, 561nm, and 647nm) were controlled via Nikon Elements software. Nikon objectives used include 20x (0.75NA), 40x (0.95NA) or 60x oil (1.4NA). Optical sectioning was performed at Nyquist for the longest wavelength. Analysis of mitochondrial length and occupancy were performed in Nikon Elements.

Live imaging-Electroporated cortical neurons were imaged at 15-21DIV with EMCCD camera (Andor, iXon3-897) on an inverted Nikon Ti-E microscope (40x objective NA0.95 with 1.5x digital zoom or 60x objective NA1.4) with Nikon Elements. 488nm and 561nm lasers shuttered by Acousto-Optic Tunable Filters (AOTF) or 395nm, 470nm, and 555nm Spectra X LED lights (Lumencor) were used for the light source, and a custom quad-band excitation/dichroic/emission cube (based off Chroma, 89400) followed by clean up filters (Chroma, ET435/26, ET525/50, ET600/50) were applied for excitation and emission. We used modified normal tyrode solution as a bath solution at 37°C (Tokai Hit Chamber), which contained (in mM): 145 NaCl, 2.5 KCl, 10 HEPES pH7.4, 2 CaCl_2_, 1 MgCl_2_, 10 glucose.

For ATEAM (gift from Hiromi Imamura) imaging a CFP/YFP cube (Chroma, 59217) followed by clean up filters (Chroma, ET475/20, ET540/21) was used, while for roGFP2 (Addgene) imaging, the custom cube above was used followed by clean up with ET525/50.

For Tetramethylrhodamine (Sigma, TMRM) imaging, cells were incubated in the solution above and 20nM TMRM for 20 minutes at 37°C, and imaged in the solution with 5nM TMRM.

For calcium imaging on evoked release, we added APV (50μM, Tocris) and CNQX (20μM, Tocris) in bath solution. Evoked releases were triggered by 1ms current injections with a concentric bipolar electrode (FHC) placed 20μm away from transfected axons. We applied 20APs at 10Hz with 30V using the stimulator (Model 2100, A-M systems) and imaged with 500ms or 100ms interval (2Hz or 10Hz) during 90sec for mt-GCaMP5G signals and 100ms interval (10Hz) for 15sec for VGLUT1-GCaMP5G signals. At the end of acquiring, we added the calcium ionophore ionomycin (5μM, EMD Millipore) and continued imaging with 1sec interval to obtain F_max_ value. For blocking mitochondrial Na^+^/Ca^2+^ exchanger, we added CGP 37157 (10μM, Tocris) following mt-GCaMP5G imaging with 500ms interval. After 3min, we imaged the same area with the identical condition (20APs at 10Hz).

For syp-pHluorin imaging, 20APs were applied at 10Hz and neurons were imaged with 100ms interval (10Hz) during 60sec, then the bath solution was changed with tyrode solution containing 55mM NH_4_Cl for F_max_ value.

Images were analyzed using a Fiji (Image J) plug-in, Time Series Analyzer (v3.0). Each vGLUT1-GCaMP5G or syp-pHluorin puncta and nearby backgrounds were selected by circular ROIs and intensities were measured by plug-in. For mt-GCaMP5G measurement, single synapse-associated mitochondria were pooled for comparing uptake of presynaptic Ca^2+^ from a single presynaptic bouton. Full-length mitochondria were marked by a freehand selection tool and total intensities were measured. After intensities were corrected for background subtraction, ΔF values were calculated from (F-F_0_). F_0_ values were defined by averaging 10 frames before stimulation, and F_max_ values were determined by averaging 10 frames of maximum plateau values following ionomycin or NH_4_Cl application, then, used for normalization. Long mitochondria in *Mff* knockdown axons for Ca^2+^ and pHluorin imaging were selected by the length over 3µm. Diffusion of long mitochondrial Ca^2+^ from a single presynapse is also analyzed by Fiji. Line ROIs (1µm, width 3) were placed on a presynapse-overlapped site and every 2µm distance from the site. Then, Plot Z-axis Profile function was performed for obtaining intensities from each line, and time to peak was defined from the plot.

### Cell Line Transfection and Western Blot

HEK cells were transfected via jetPrime (Polyplus) by following the manufacture‘s instructions. Cells were harvested in cold DPBS and lysed in ice-cold lysis buffer containing 25 mM Tris (pH7.5), 2 mM MgCl_2_, 600 mM NaCl, 2 mM EDTA, 0.5% NP-40, 1X protease and phosphatase cocktail inhibitors (Sigma) and Benzonase (0.25 U/μl of lysis buffer; Novagen). Aliquots of the proteins were separated by SDS-PAGE and then transferred to a polyvinylidene difluoride (PVDF) membrane (Amersham). After transfer, the membrane was washed 3X in Tris Buffer Saline (10 mM Tris-HCl pH 7.4, 150 mM NaCl) with 0.1% of Tween 20 (T-TBS), blocked for 1 hr at room temperature in Odyssey Blocking Buffer (TBS, LI-COR), followed by 4°C overnight incubation with the appropriate primary antibody in the above buffer. The following day, the membrane was washed 3X in T-TBS, incubated at room temperature for 1 hr with IRDye secondary antibodies (LI-COR) at 1:10,000 dilution in Odyssey Blocking Buffer (TBS), followed by 3X T-TBS washes. Visualization was performed by quantitative fluorescence using an Odyssey CLx imager (LI-COR). Signal intensity was quantified using Image Studio software (LI-COR). Primary antibodies used for Western-blotting are mouse anti-HA (1:2,000, Covance), rabbit anti-ERp72 (1:1000, Cell Signaling Technologies).

### Quantification and Statistical Analysis

All statistical analysis and graphs were performed/created in Graphpad’s Prism 6. Statistical tests, p values, and (n) numbers are presented in the figure legends. Gaussian distribution was tested using D’Agostino & Pearson’s omnibus normality test. We applied non-parametric tests when data from groups tested deviated significantly from normality. All analysis were performed on raw imaging data without any adjustments. Images in figures have been adjusted for brightness and contrast (identical for control and experimental conditions in groups compared).

### RESOURCES

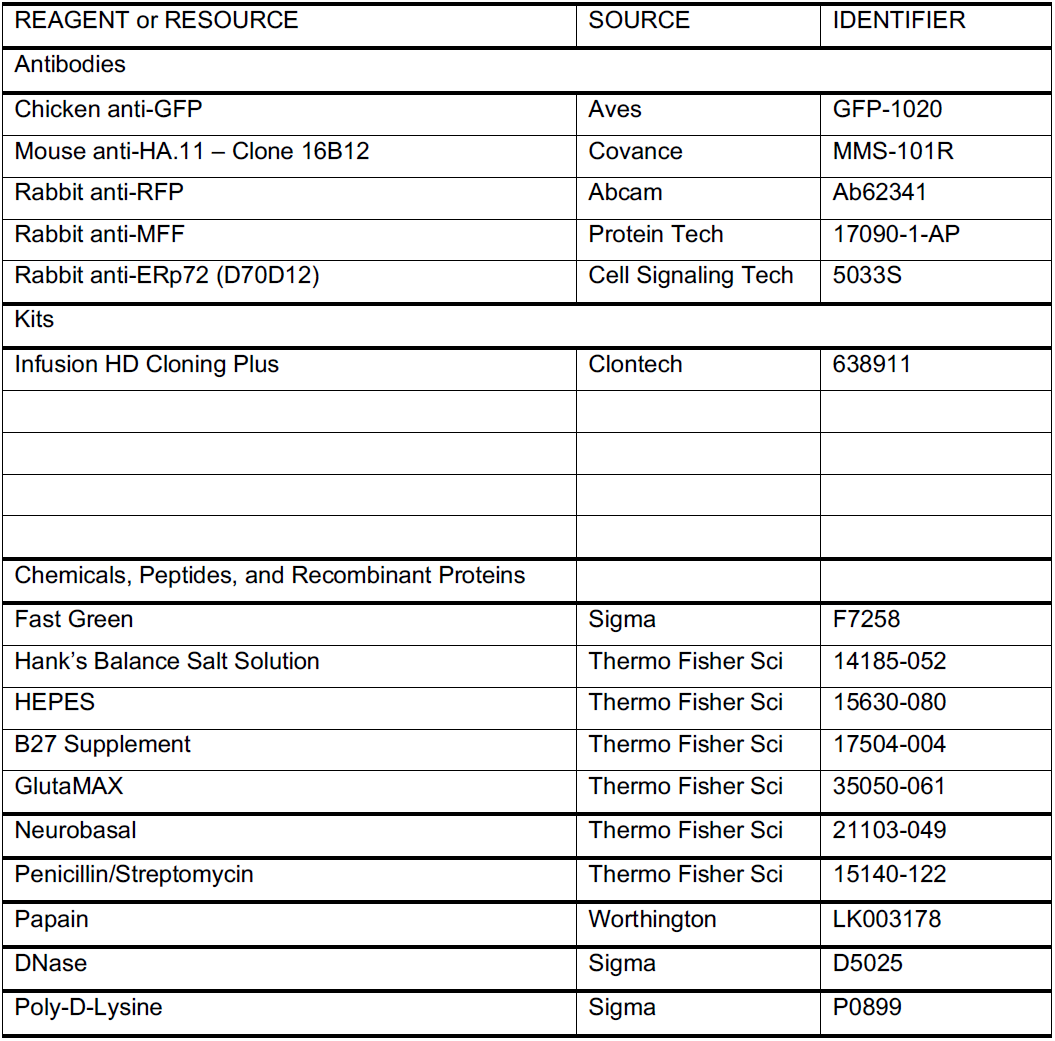
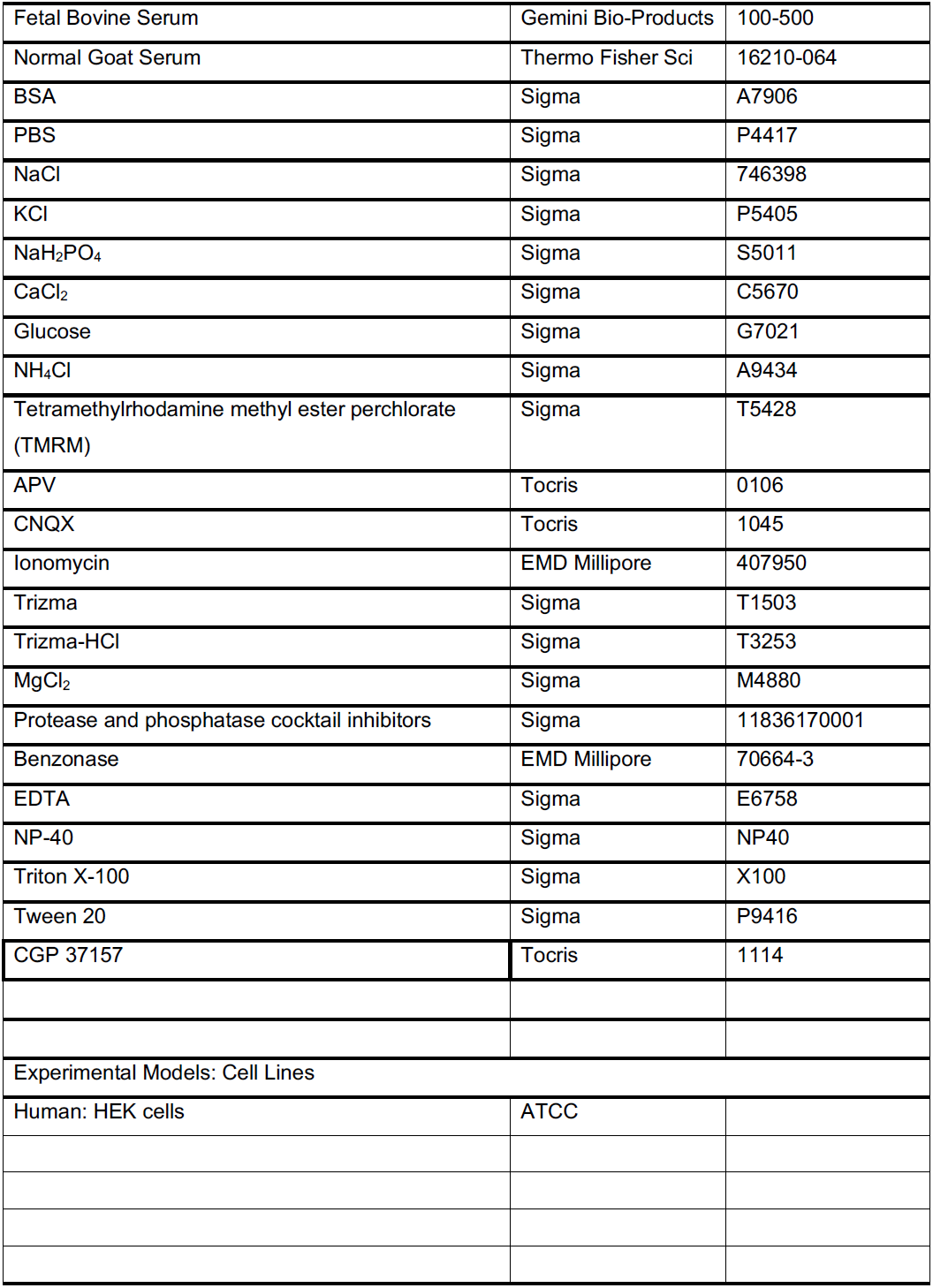
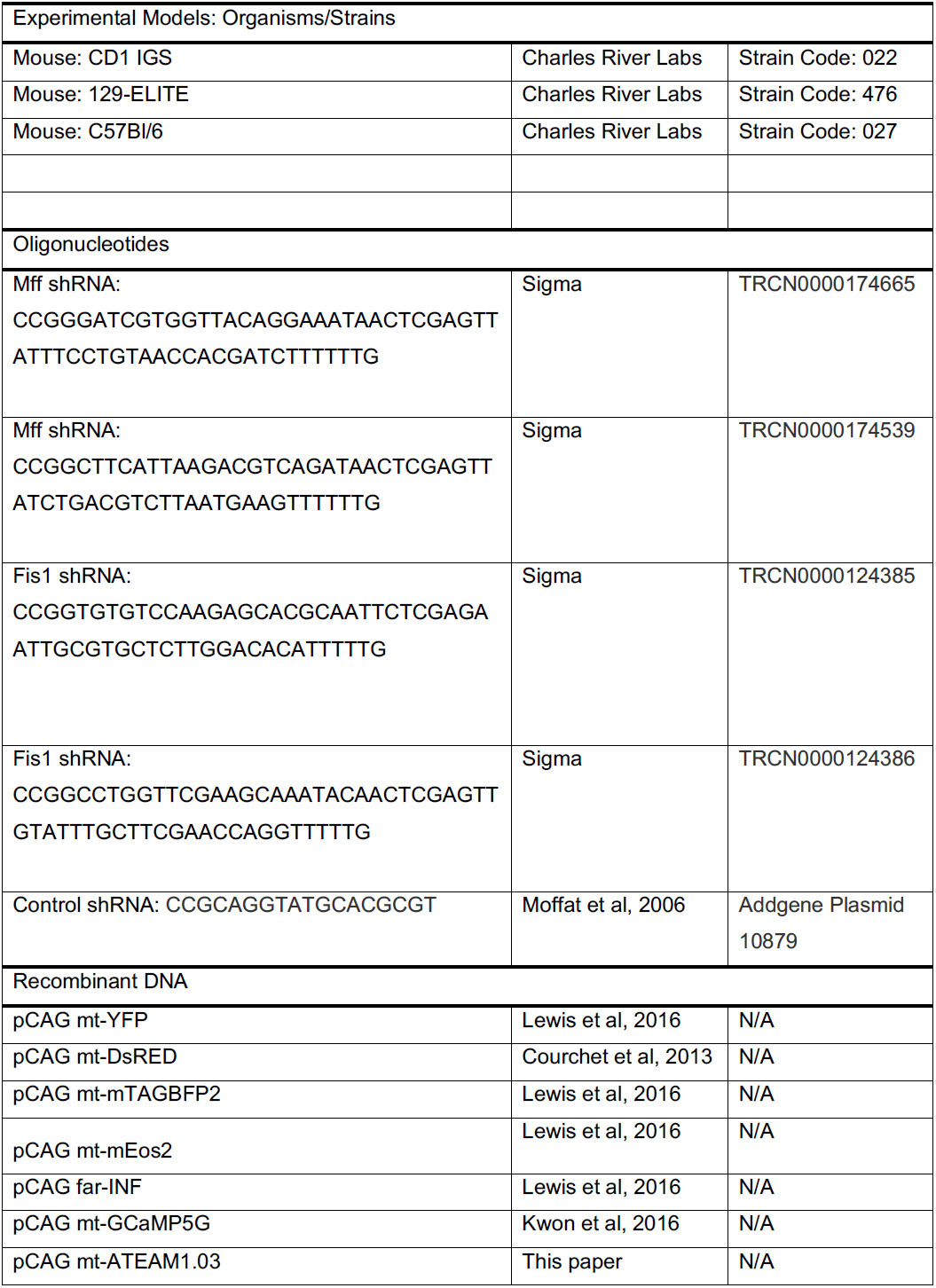
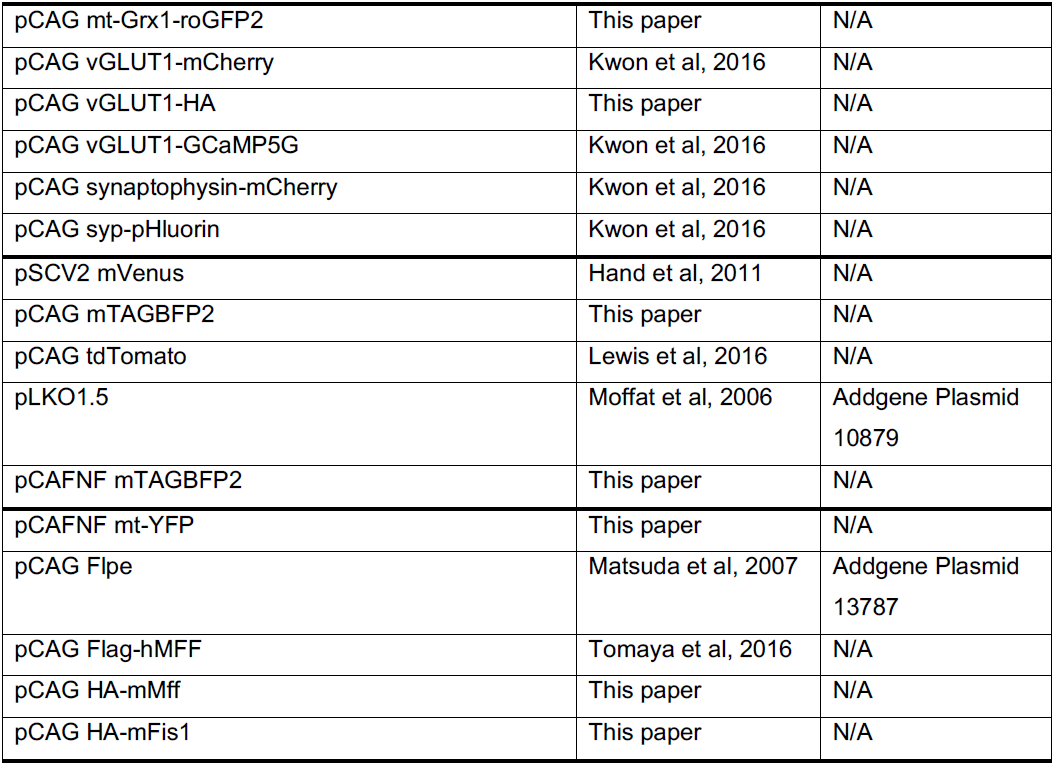

**Supplemental Figure 1. Validation of *Mff* shRNA knockdown constructs**

(**a**) Western blot of shRNA knockdown efficiency via overexpression of HA-tagged mouse MFF (top) in HEK cells. Antibody against an endogenous ER protein (ERp72; bottom) was used as the loading control. (**b**) Western blot of shRNA knockdown efficiency via overexpression of HA-tagged mouse Fis1 (top) in HEK cells. Endogenous ERp72 (bottom) was used as the loading control. (**c-e**) Representative images of a neuron electroporated with mt-YFP and control shRNA via *ex utero* electroporation at E15.5, and stained at 7DIV with antibodies for GFP and Mff. (**f-h**) Representative images of neurons electroporated with mt-YFP and a (1:1) mixture of *Mff* shRNAs via *ex utero* electroporation at E15.5, and stained at 7DIV with antibodies for GFP and Mff. (**i-k**) Representative images of an axon from a neuron electroporated with mt-YFP and control shRNA via *ex utero* electroporation at E15.5, and stained at 7DIV with antibodies for GFP and MFF. (**l-n**) Representative images of an axon from a neuron electroporated with mt-YFP and a (1:1) mixture of *Mff* shRNAs via *ex utero* electroporation at E15.5, and stained at 7DIV with antibodies for GFP and Mff. Orange arrows point to the same mitochondria in each set of images. Related to **Figure 1.**

**Supplemental Figure 2. Decreased MFF activity does not increase dendritic mitochondrial length**

(**a-c**) Representative images of a dendrite from a neuron electroporated with mt-YFP, tdTomato and control shRNA via *in utero* electroporation at E15.5, and stained at P21 with antibodies for GFP and tdTomato. (**d-f**) Representative images of a dendrite from a neuron electroporated with mt-YFP, tdTomato and a mixture of *Mff* shRNAs via *in utero* electroporation at E15.5, and stained at P21 with antibodies for GFP and tdTomato. (**g**) Quantification of mitochondrial length in the dendrites. Data is represented at minimum to maximum box plots, with the box denoting 25^th^, 50^th^ and 75^th^ percentile. (**h**) Quantification of the percent of the dendrite occupied by mitochondria. Data is represented at minimum to maximum box plots, with the box denoting 25^th^, 50^th^ and 75^th^ percentile. n_control shRNA_ = 26 dendrites, 267 mitochondria; n_MFF shRNA_ = 19 dendrites, 146 mitochondria. n.s., p>0.05 according to Mann-Whitney test. Related to **Figure 2.**

**Supplemental Figure 3. Fewer mitochondria enter the axon but have increased length upon loss of MFF expression**

(**a**) Schematic of the imaging paradigm used for measuring axonal entry of mitochondria via mitochondrial-targeted mEos2. (**b**) Selected timeframes of mitochondria entering the axon of a 10DIV neuron *ex utero* electroporated with control shRNA and mt-mEos2. (**c**) Selected timeframes of mitochondria entering the axon of a 10DIV neuron *ex utero* electroporated with *Mff* shRNA and mt-mEos2. Fewer mitochondria enter the axon upon *Mff* knockdown, but are much longer. See supplemental video 2. (**d**) Quantification of the number of axonal entry events per hour in 7-11DIV axons. Data is represented as a scatter plot with mean ± sem. ***p<0.001 for control vs. *Mff* shRNA. Mann-Whitney test. (**e**) Cumulative frequency of mitochondrial length upon axonal entry for control and *Mff* shRNA mediated knockdown in 7-11DIV axons. **** p<0.0001 for control vs. *Mff* shRNA fission. Mann-Whitney test. n_control shRNA_ = 12 axons, 63 mitochondria; n_MFF shRNA_ = 19 axons, 48 mitochondria. Related to **Figure 3.**

**Supplemental Figure 4. Loss of MFF activity does not affect the final localization of axonal mitochondria at presynaptic sites**

(**a**) Quantification of the percent of stationary mitochondria for 15min at 7DIV. Data is represented as a scatter plot with mean ± sd. n_control shRNA_ = 14 axons; n_MFF shRNA_ = 16 axons. See supplemental video 3. (**b**) Quantification of the percent of stationary mitochondria for 15min at 21DIV. Data is represented as a scatter plot with mean ± sd. n_control shRNA_ = 10 axons; n_MFF shRNA_ = 9 axons. (**c**) Quantification of average velocity for motile mitochondria in axons from (A) Data is represented as a scatter plot with mean ± sd. (**d**) Quantification of run length for motile mitochondria in axons from (A). Data is represented as a scatter plot with mean ± sd. (**e**) Representative images of an axon from a neuron electroporated with mt-YFP, VGLUT1-HA, tdTomato and control shRNA (left panels) or a mixture of *Mff* shRNAs (right panels) via *in utero* electroporation at E15.5, and stained at P21 with antibodies for GFP, HA and tdTomato. (**f**) Quantification of the percent of mitochondria at presynaptic sites. (**g**) Quantification of the percent of presynaptic sites with mitochondria. (**h**) Quantification of the number of VGLUT1-HA puncta per 100 microns of axon. Data is represented as scatter plots with mean ± sd. n_control shRNA_ = 16 axons; n_MFF shRNA_ = 26 axons. Mann-Whitney test; p values in figure. Related to **Figures 5 and 6.**

**Supplemental Figure 5. Paradigm for the measure of mitochondrial membrane potential in both control and *Mff***-deficient dissociated neuronal co-culture

(**a**) Schematic representation of how neurons were labeled with color-swapped fluorescent proteins to identify axons from control and *Mff* shRNA-expressing neurons before TMRM labeling. Following *ex utero* electroporation with DNA plasmids encoding control shRNA, mt-mTAGBFP2 and YFP or *Mff* shRNA, mt-YFP and mTAGBFP2, dissociated neurons were mixed at a 1:1 ratio and plated on coverslips for co-cultures. This strategy allows for the simultaneous imaging of mitochondrial membrane potential (TMRM-see **Fig. 5a-b**) in both control and *Mff* knockdown neurons under the exact same culture conditions. (**b**) Representative field of view for mTAGBFP2/mt-mTAGBFP2. (**c**) Representative field of view for YFP/mt-YFP. (**d**) Merged channels shown in B-C where control (sky blue arrowheads) and *Mff* knockdown (orange arrows) axons can be visualized in the same field of view. Related to **Figure 5.**

**Supplemental Figure 6. Long mitochondria in *Mff* knockdown neurons show diffusion of imported Ca^2+^**

(**a-b**) Cropped mitochondrial Ca^2+^ time-lapse images of control and *Mff* knockdown axons. Presynaptic mitochondrial Ca^2+^ signals were captured with 100ms interval for monitoring diffusion of imported Ca^2+^ through mitochondrial matrix. Ca^2+^ propagation in long mitochondria of *Mff* knockdown axons occurs from presynaptic sites. (**c**) For quantification of diffusion time, analysis was performed with single presynapse-overlapped mitochondria from *Mff* knockdown neurons. (**d-e**) Graphs display the latency of time to peak depending on distance from presynaptic sites. n_MFF shRNA_ = 8 dishes, 8 mitochondria. p=0.0012 for 0 vs. 4, 0.0067 for 0 vs. 6. One-way ANOVA, Bonferroni‘s multiple comparison test. Related to **Figure 6.**

**Supplemental Figure 7. MFF-dependent mitochondrial fission regulates presynaptic release and axon branching by limiting axonal mitochondria size**

(**a**) In control neurons, MFF activity is required for the small size of axonal mitochondria both upon axonal entry, as well as for maintenance along the axon. Mitochondrial size is maintained by coupling the majority of mitochondrial fusion events to a fission event. (**b**) Loss of MFF activity increases mitochondrial entry size and decreases the ratio of fission to fusion along the axon. This leads to increased mitochondrial size along the axon and reduced terminal branching. (**c**) In control axons, ~50% of presynaptic boutons are occupied by mitochondria. Upon neuronal activity, the mitochondria buffer a significant amount of Ca^2+^ influx, in an MCU dependent manner, thereby reducing presynaptic vesicle release as compared to presynaptic sites without a mitochondria. (**d**) Upon MFF knockdown, the increased mitochondrial matrix volume allows for increased Ca^2+^ uptake after neuronal activity, and strongly reduces presynaptic vesicle release at these sites. The reduction in terminal axon branching observed following MFF knockdown is likely due to this reduction in presynaptic release as presynaptic release is required for the stabilization of axonal branches.

**Supplemental Movie 1: Mitochondrial fission and fusion dynamics along the axon upon *Mff* knockdown. Related to Figures 2 & 3.**

**Supplemental Movie 2: Dynamics of mitochondrial entry into the axon upon *Mff* knockdown. Related to Figures 2 and 3.**

**Supplemental Movie 3: Axonal mitochondrial motility upon *Mff* knockdown. Related to Figures 2 & 3.**

**Supplemental Movie 4: Presynaptic mitochondrial Ca^2+^ uptake upon evoked activity in *Mff* knockdown axons, Related to Figure 6.**

